## Supplemental Figures S1 – S23 for "FOXJ1 mediates taxane resistance through regulation of microtubule dynamics"

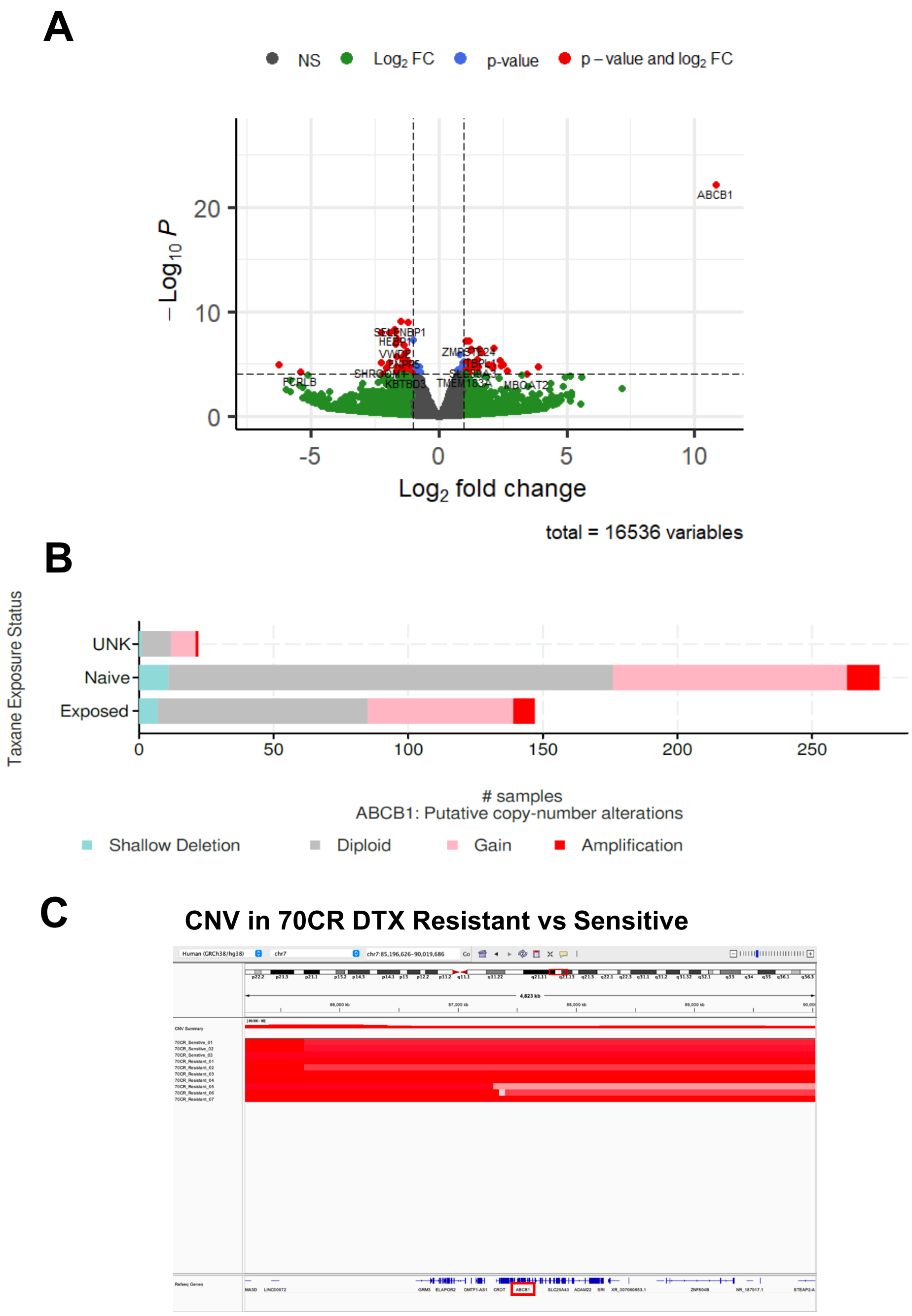


**Figure S1. ABCB1 overexpression and amplification in docetaxel resistant LuCAP 70CR PDXs. (A)** Differential gene expression for LuCaP 70CR resistant versus sensitive PDXs. **(B)** ABCB1 CNV in PC that were taxane exposed (5.44%) versus naïve (4.36%) or unknown (UNK). Data are from the SU2C PC Dream Team and analyzed on CBioPortal. (**C**) Copy number variation in LuCaP 70CR resistant and sensitive PDXs visualized by Integrative Genomics Viewer (IGV).

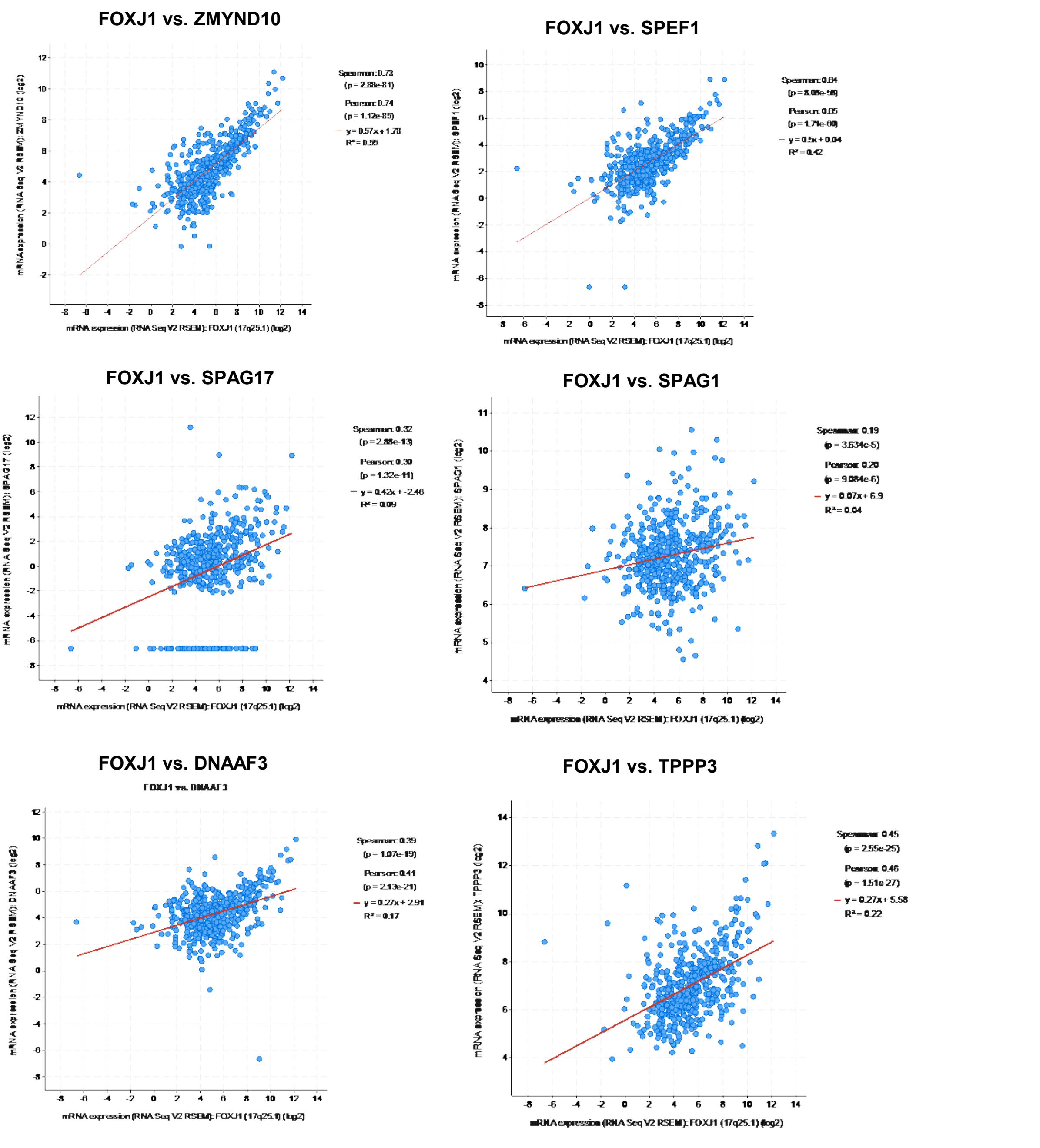


**Figure S2. FOXJ1 co-expression with cilium-related genes in primary PC.** Expression of FOXJ1 is plotted versus indicated cilium related genes in TCGA Firehose Legacy dataset.

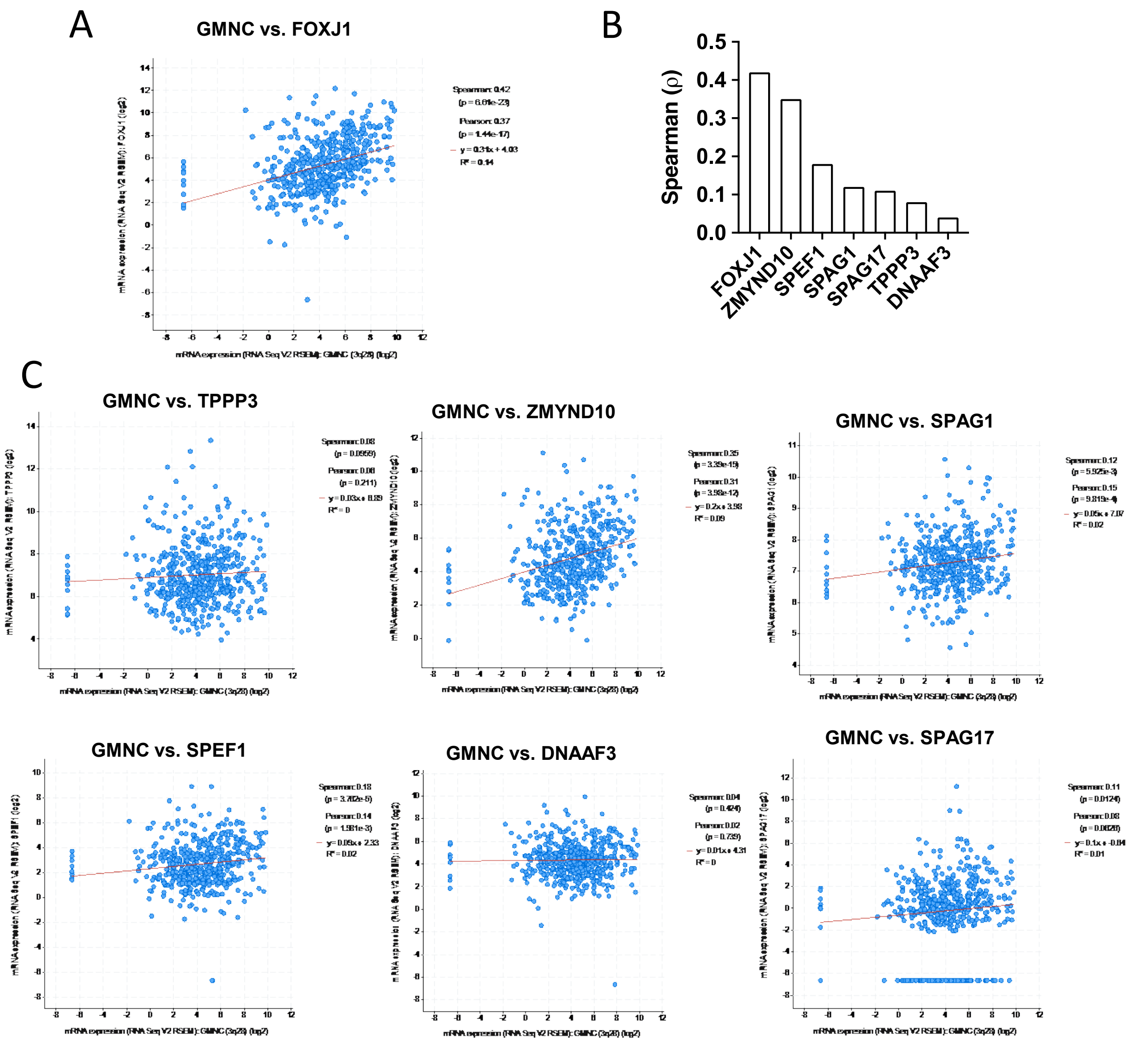


**Figure S3. GEMC1 co-expression with FOXJ1 and cilium-related genes in TCGA Firehose Legacy primary PC dataset. (A)** GEMC1 versus FOXJ1 expression. **(B)** Summary of correlations between GEMC1 and indicated cilium-related genes in TCGA Firehouse Legacy PC dataset. **(C)** Plots of GEMC1 versus cilium-related genes.

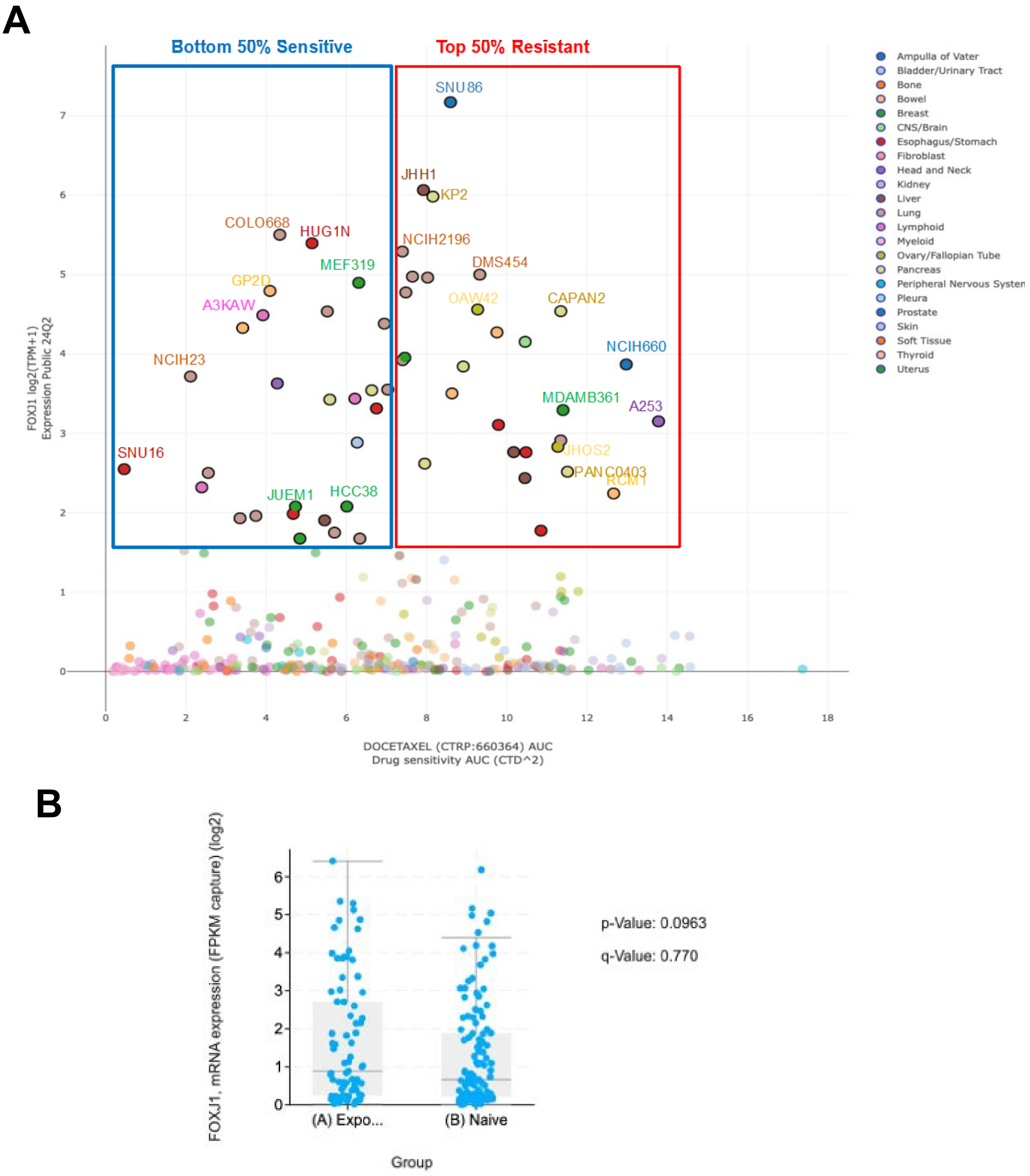


**Figure S4. Increased FOXJ1 is associated with cell line docetaxel-resistance and drug exposure in patients. (A)** Response to docetaxel treatment in DepMap's 300 cancer cells. Each dot represents one individual cancer cell line. The cell lines with top FOXJ1 expression (>1.5) were selected to be compared based on the AUC scores (Top50% vs Bottom50%) in **Figure 1G**. **(B)** FOXJ1 mRNA expression in taxane exposed and naïve patients from SU2C data on cBioPortal.

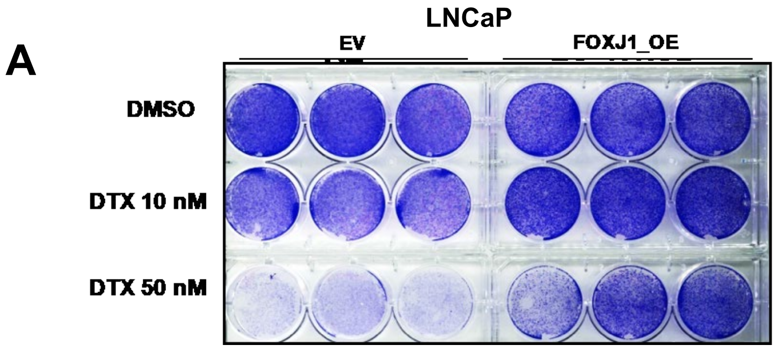


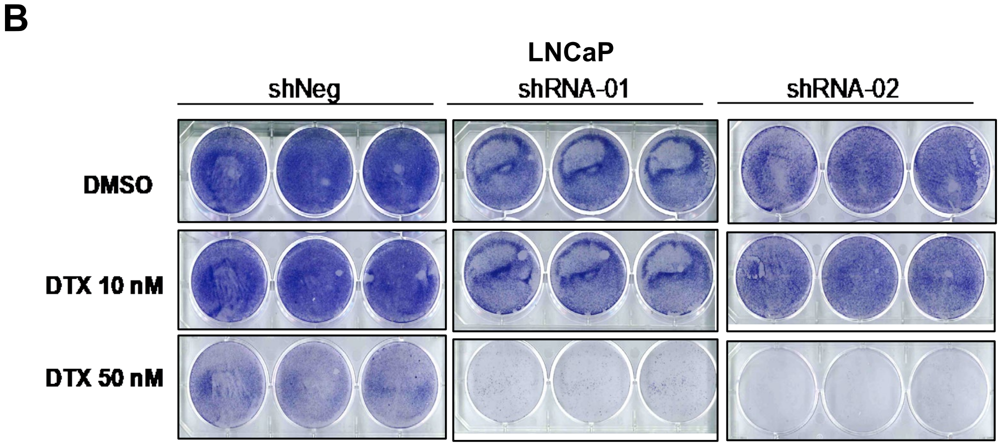


**Figure S5. FOXJ1 effects on colony formation in response to docetaxel. (A)** Images for colony formation in LNCaP EV and LNCaP FOXJ1_OE cells with indicated treatments. **(B)** Images for colony formation in LNCaP cells expressing negative control shRNA (shNeg) or FOXJ targeted shRNA (shRNA-01 and shRNA-02) with indicated treatments.

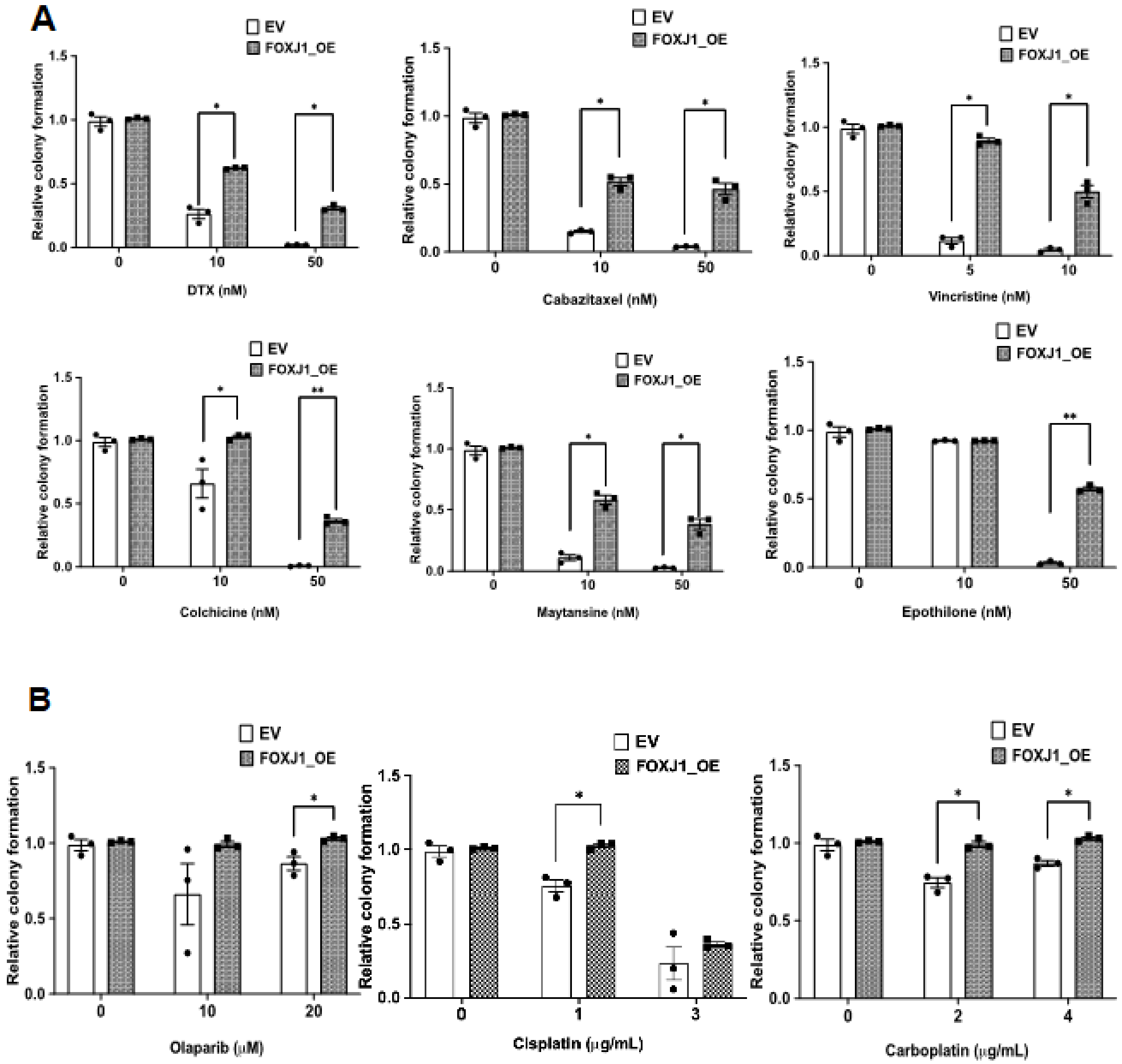


**Figure S6. FOXJ1 overexpression decreases sensitivity to additional tubulin target drugs and additional agents.** (A) LNCaP cells overexpressing FOXJ1 or empty vector (EV) control were plated at high density and treated for 7 days with the indicated drugs. The drugs were then washed out and cells were cultured for an additional 1-3 weeks until colonies were visible. Colonies were then stained with crystal violet and quantified. (B) Cells as in (A) were treated as indicated.

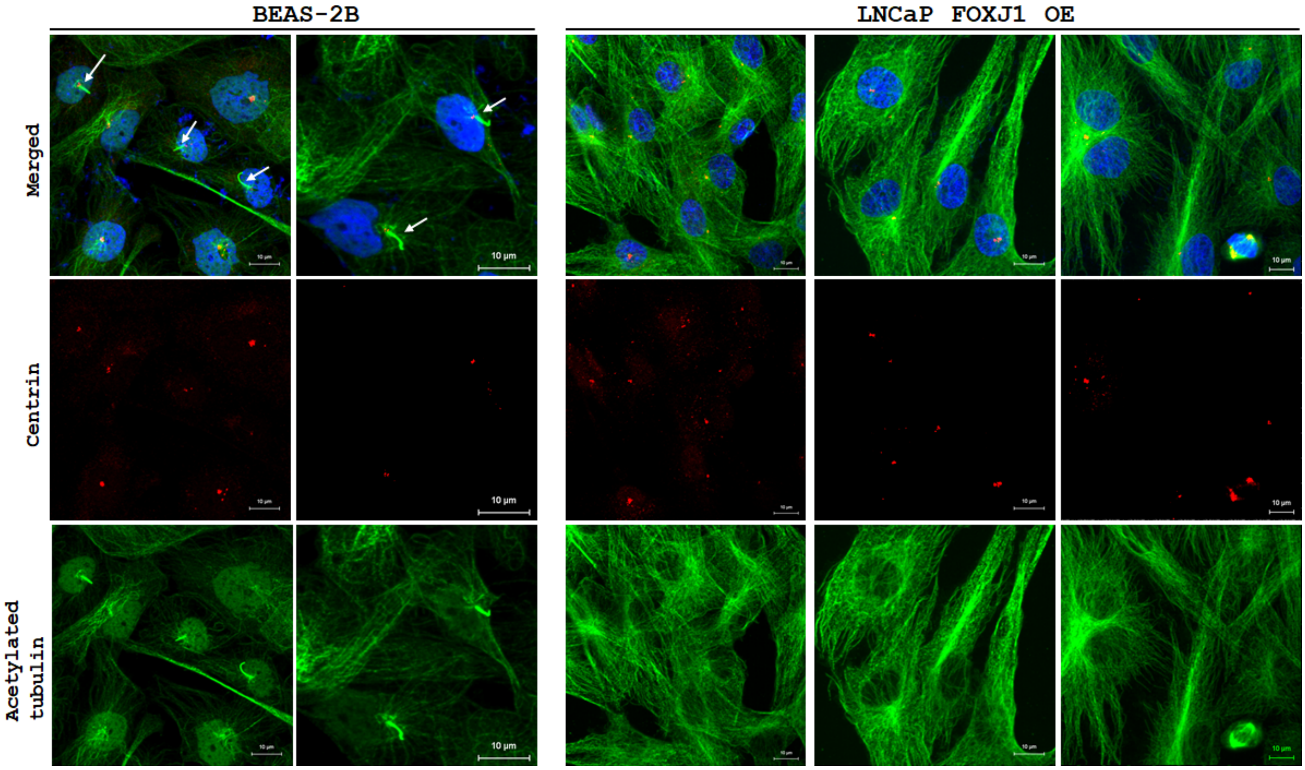


**Figure S7. Assessment of Cilia Formation via Acetylated α-Tubulin and Centrin Staining.** Representative microscopy images are shown for BEAS-2B cells (left 2 panels) and LNCaP FOXJ1-overexpressing (OE) cells (right 3 panels). The top images display merged fluorescence for acetylated α-tubulin (green), centrin (red), and DAPI (blue). In BEAS-2B cells, cilia are clearly visible, as indicated by the arrows. The corresponding single-channel images are shown below: centrin in red at the ciliary base and acetylated α-tubulinin in green marking the cilia. In contrast, LNCaP FOXJ1 OE cells lacked detectable cilia under the same staining conditions, as evidenced by the absence of acetylated α-tubulin-positive projections.

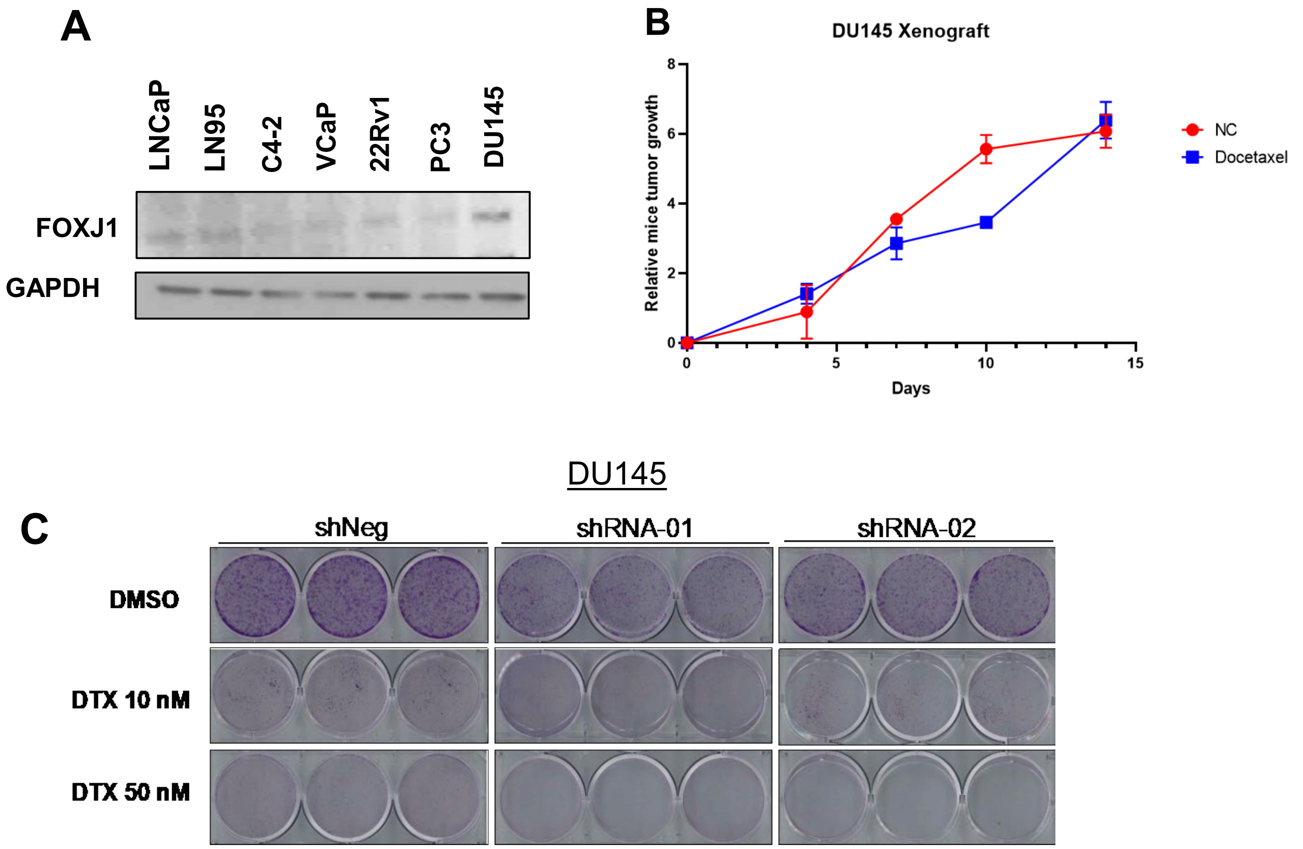


**Figure S8. FOXJ1 knockdown sensitizes DU145 cells to docetaxel. (A)** FOXJ1 protein expression in multiple PC cell lines. **(B)** DU145 xenografts were allowed to grow until ~250 $mm^3$ and were then treated with 30 mg/kg docetaxel at Day 0, 5 mice for each group. Tumor growth was then monitored for two weeks. **(C)** Images for colony formation in DU145 shNeg and DU145 FOXJ1 knockdown (shRNA-01 and shRNA-02) cells with indicated docetaxel treatments.

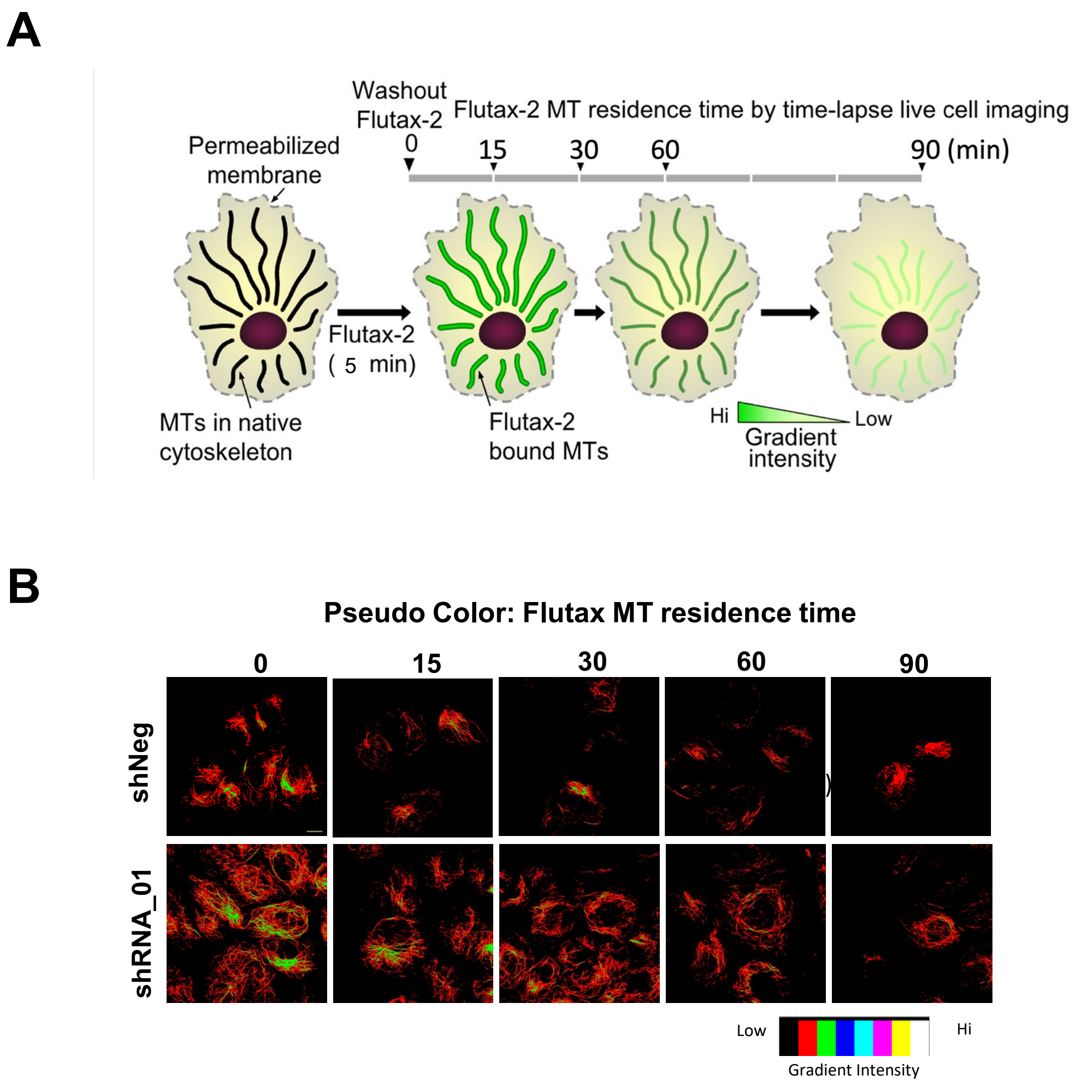


**Figure S9. FOXJ1 knockdown increases Flutax residence time on microtubules. (A)** Experimental design: Native cytoskeletons from DU145 shNeg and shRNA_01 cells were treated with 1 μM FITC-conjugated paclitaxel (Flutax-2) for 5 minutes. The residence time of Flutax-2 on microtubules was tracked using time-lapse live cell imaging for 90 minutes post-washout with a Zeiss spinning disk microscope. **(B)** Pseudo-color representations of Flutax-2 intensity levels on cellular microtubules. The red to yellow color gradient represents low to high intensities, respectively.

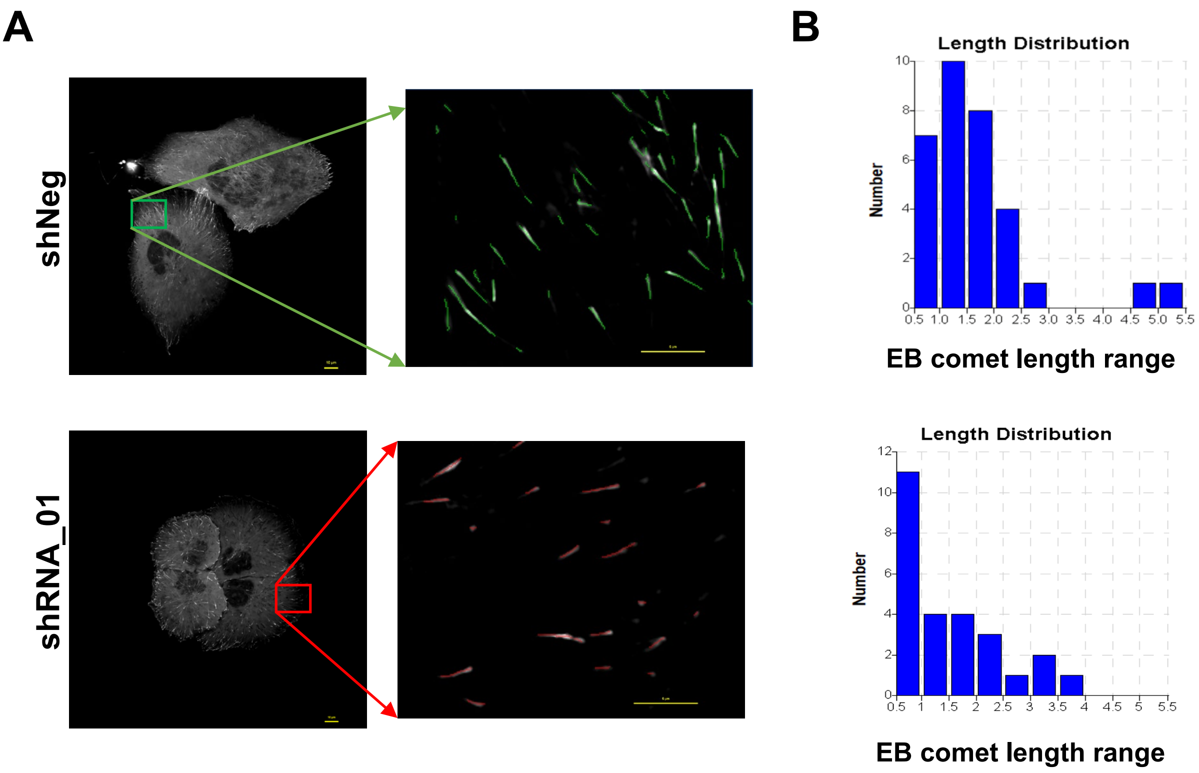


**Figure 10. FOXJ1 knockdown significantly affects EB1 comet length. (A)** Representative grayscale images of EB1-EGFP comets in DU145 shNeg and shRNA_01 cells are shown, with each image featuring a scale bar of 10 μm. The highlighted square within each image delineates the leading-edge area chosen for zoomed-in visualization. The zoomed-in images provide a closer view of EB1-EGFP comet length (computer-generated) in a single Z-plane, with a scale bar of 5 μm. **(B)** The bar graph illustrates the differential length distribution between DU145 shNeg and shRNA_01 cells, as observed in panel (A). This distribution was quantified using Nikon Element software across length intervals. The X-axis represents length range bins from 0.5-1 μm up to 5-5.5 μm, while the Y-axis indicates the number of EB1 comets in each length range.

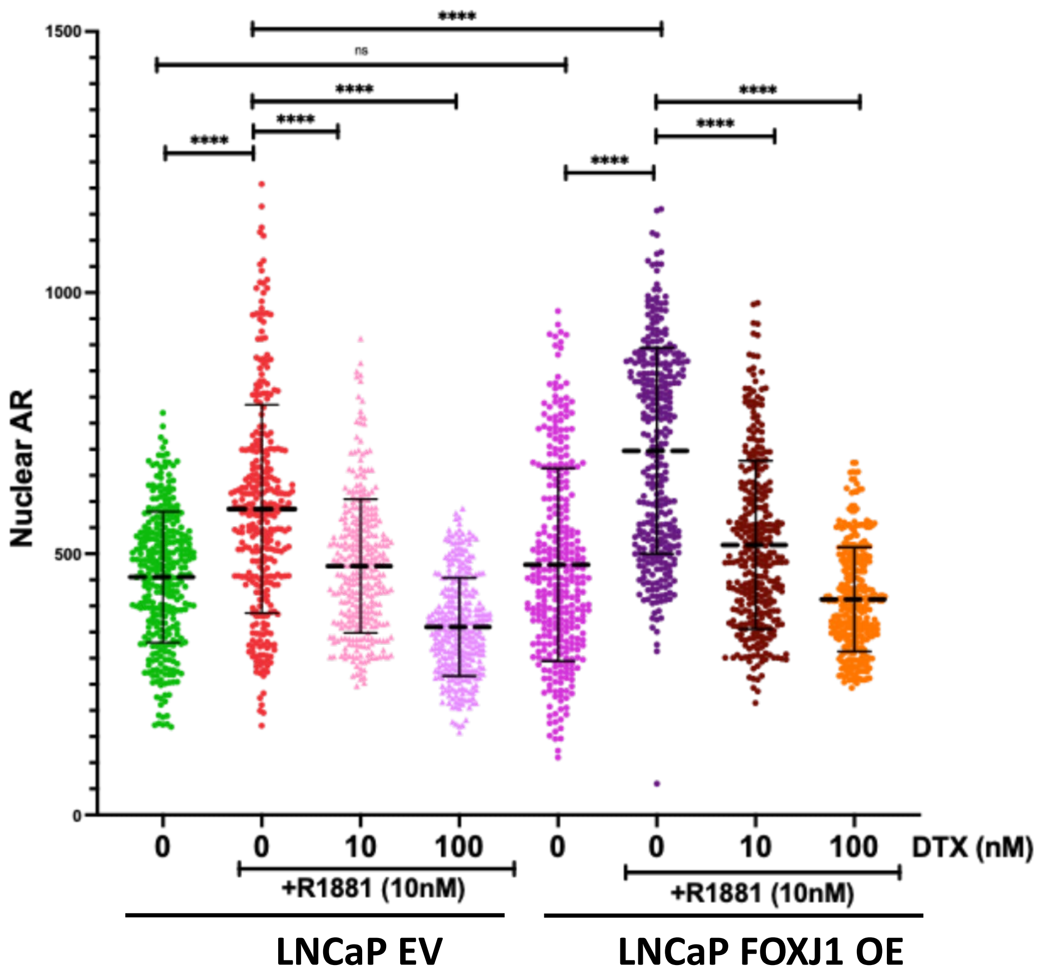


**Figure S11. FOXJ1 overexpression increases AR nuclear localization in response to androgen.** Quantification of nuclear AR levels from immunofluorescence images. Data are presented as mean ± SD. One-Way ANOVA was used to compare AR levels within each cell line's drug treatment groups. Independent samples t-tests were performed for comparisons between the LNCaP parental and LNCaP FOXJ1 OE cell lines. In both LNCaP parental and LNCaP FOXJ1 OE cell lines, R1881 treatment significantly increased nuclear AR levels (due to nuclear accumulation) compared to untreated controls. Subsequent co-treatment with DTX (10 nM and 100 nM) significantly reduced the R1881-induced nuclear AR in both cell lines, with the 100 nM DTX dose showing a stronger suppressive effect. While no significant difference was observed in nuclear AR between untreated LNCaP parental and LNCaP FOXJ1 OE cell lines, LNCaP FOXJ1 OE cells treated with R1881 exhibited significantly higher nuclear AR levels compared to LNCaP parental cells treated with R1881.

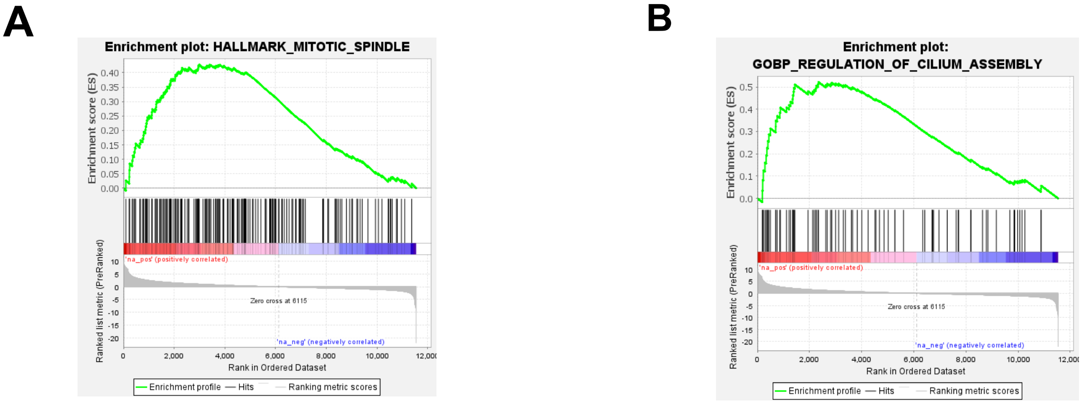


**NES 1.41**
**FDR q-value 0.16**

**NES 1.59**
**FDR q-value 0.59**

**Figure S12. GSEA for FOXJ1_OE xenografts. (A)** Enrichment for Hallmark Mitotic Spindle gene set in untreated FOXJ1_OE versus EV xenografts. **(B)** Enrichment for GOBP gene set Cilium Assembly in untreated FOXJ1_OE versus EV xenografts.

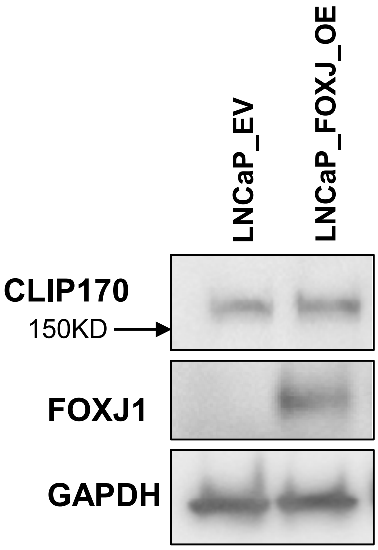


**Figure S13. CLIP170 short isoform is not increased by FOXJ1 overexpression.** LNCaP EV and FOXJ1_OE cell lines were immunoblotted for CLIP170. There was no detectable expression of the short (150 kD) isoform.

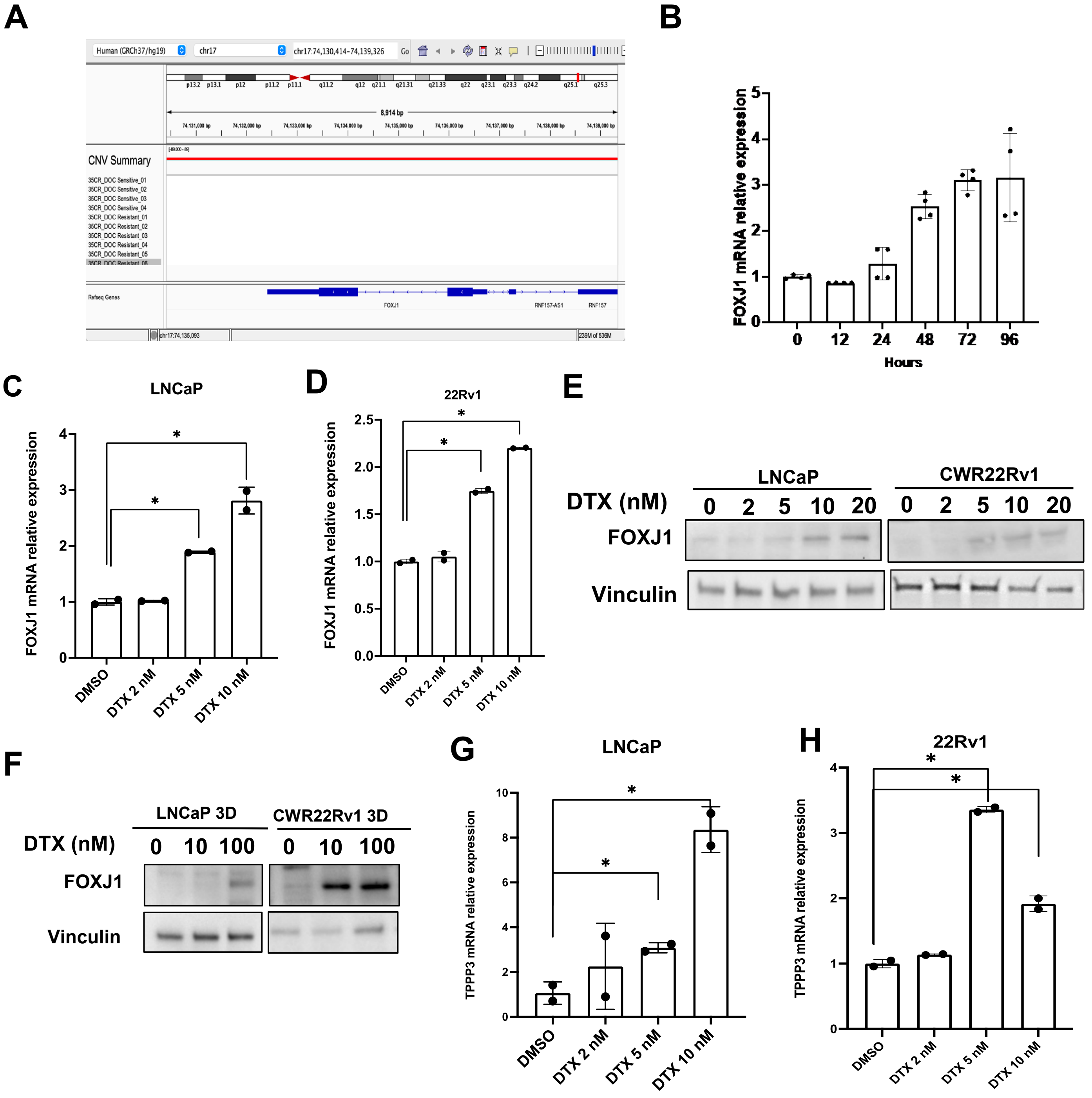


**Figure S14. FOXJ1 expression increases early in response to DTX. A,** LP-WGS shows no FOXJ1 copy number gain in LuCaP 35CR resistant PDXs as visualized on IGV. **B,** FOXJ1 mRNA expression at the indicated time points in LNCaP cells after being treated with 10 nM DTX. **C, D,** FOXJ1 mRNA expression at the indicated concentration of DTX treatment for 96 hours in LNCaP cells (C) 22Rv1 cells (D). **E,** FOXJ1 protein expression in LNCaP and 22Rv1 treated with indicated concentrations of DTX for 96 hours. **F,** FOXJ1 protein expression in 3D LNCaP and 3D 22Rv1 cultures treated with indicated concentrations of DTX for 96 hours. **G, H** TPPP3 mRNA expression in LNCaP (G) and 22Rv1 (H) cells as indicated for 96 hours.

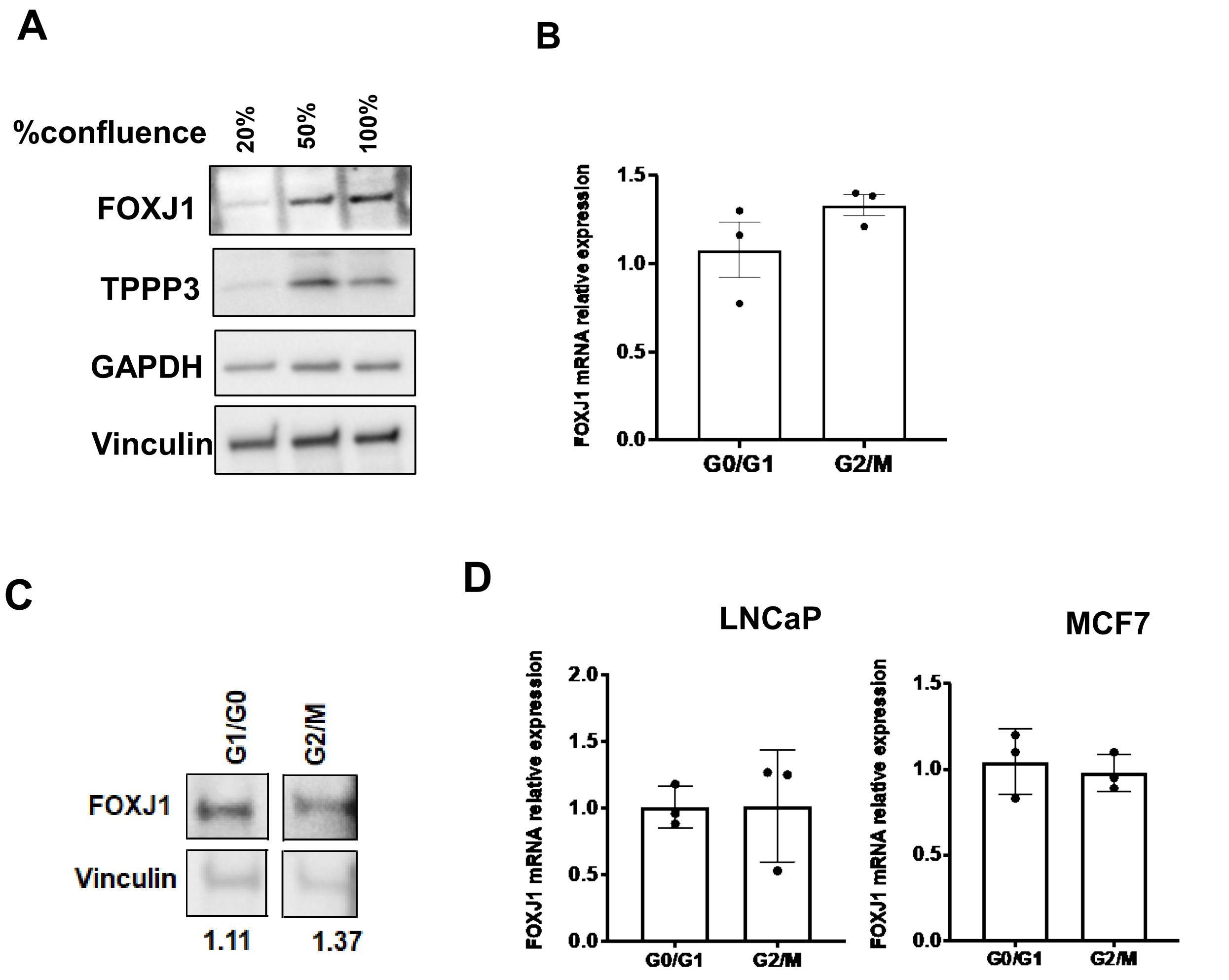


**Figure S15. FOXJ1 expression is not cell cycle regulated. A,** FOXJ1 and TPPP3 expression at different confluence levels in LNCaP cells. **B,** DU145 cells were sorted by FACS based on cell cycle. FOXJ mRNA expression in G0/G1 and G2/M populations were then determined by qRT-PCR. Levels in each were corrected based on GAPDH and the results are normalized to G0/G1. **C,** FOXJ1 protein in DU145 in G0/G1 and G2/M, ratio to vinculin is indicated. **D,** LNCaP and MCF7 cells were separated into G1/G0 and G2/M fractions by FACS. FOXJ1 mRNA levels were then determined by qRT-PCR.

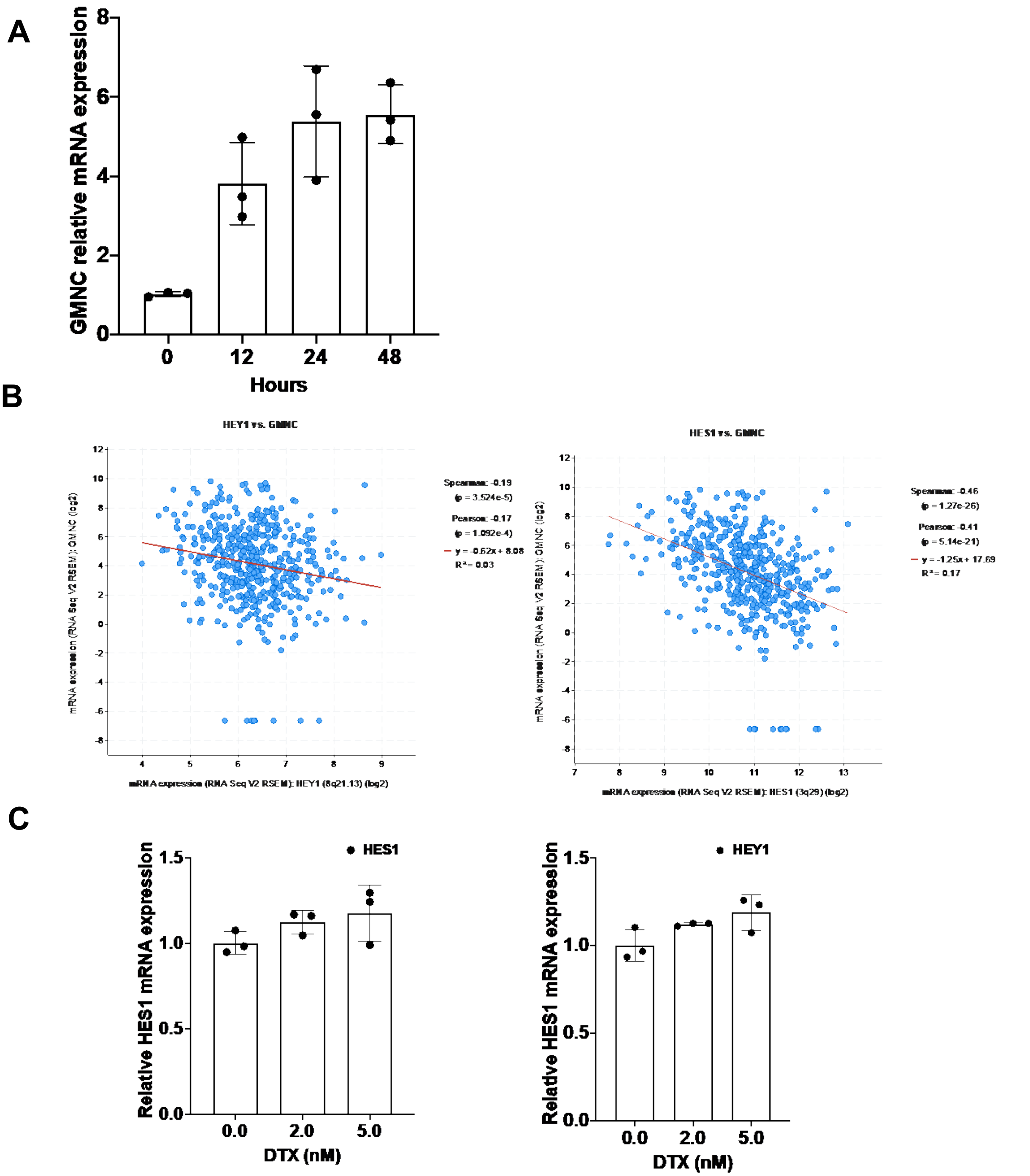


**Figure S16. GEMC1 expression is rapidly induced by DTX. A,** GEMC1 mRNA expression at the indicated time points up in LNCaP cells after being treated with 10 nM DTX. **B,** Negative correlation between expression of GEMC1 mRNA with HEY1 or HES1 mRNA in PC patients from TCGA Firehose Legacy dataset in cBioPortal. **C,** HES1 or HEY1 mRNA expression in LNCaP with indicated treatment after 24 hours.

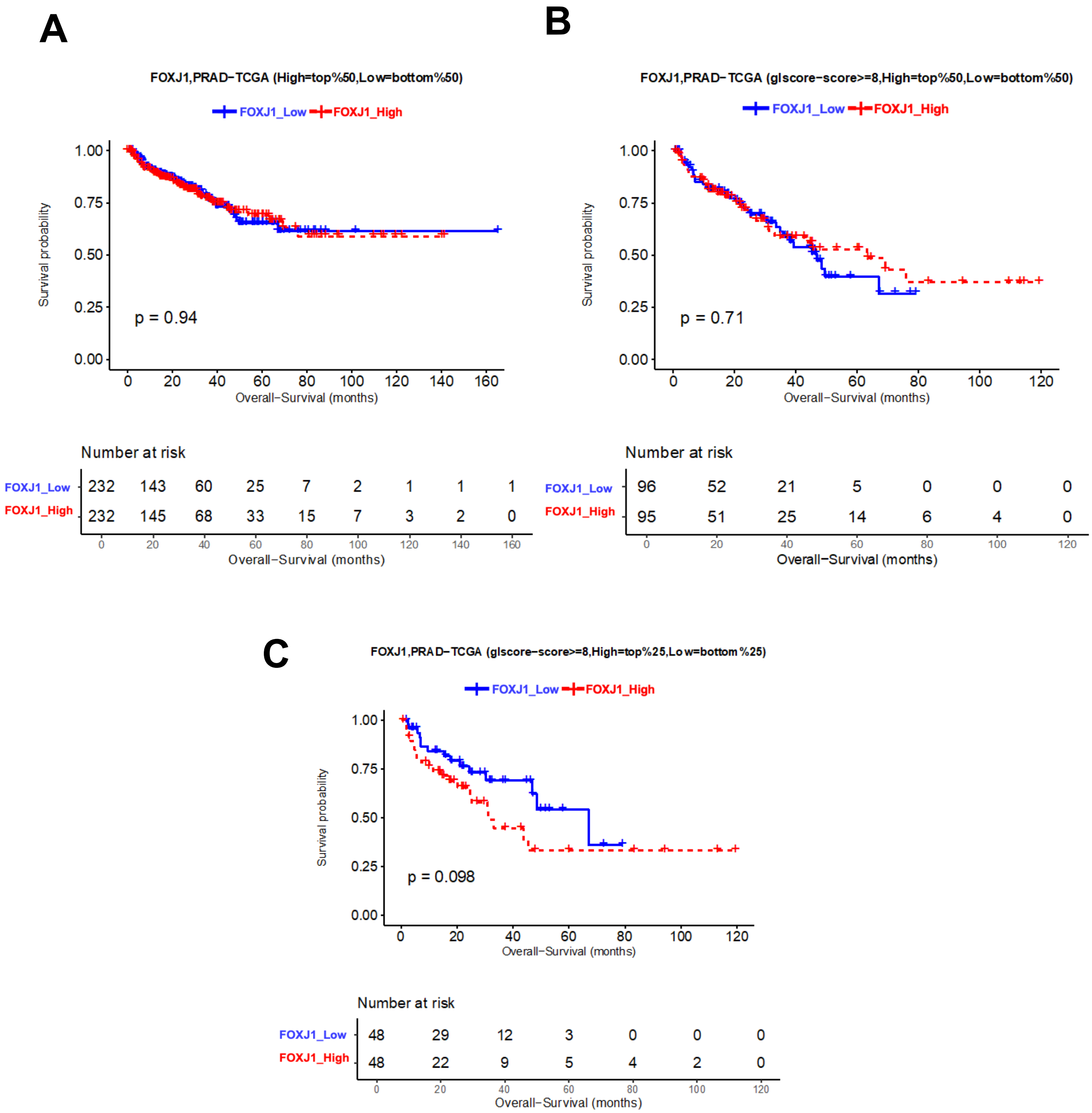


**Figure S17 Kaplan-Meier curves illustrating OS in patients from the TCGA PRAD dataset with high versus low level of FOXJ1. (A)** OS in top versus bottom 50% FOXJ1. **(B)** OS in top versus bottom 50% FOXJ1 in tumors with Gleason 8 or above. **(C)** OS in upper versus lower quartile FOXJ1 expression in tumors with Gleason 8 or above.

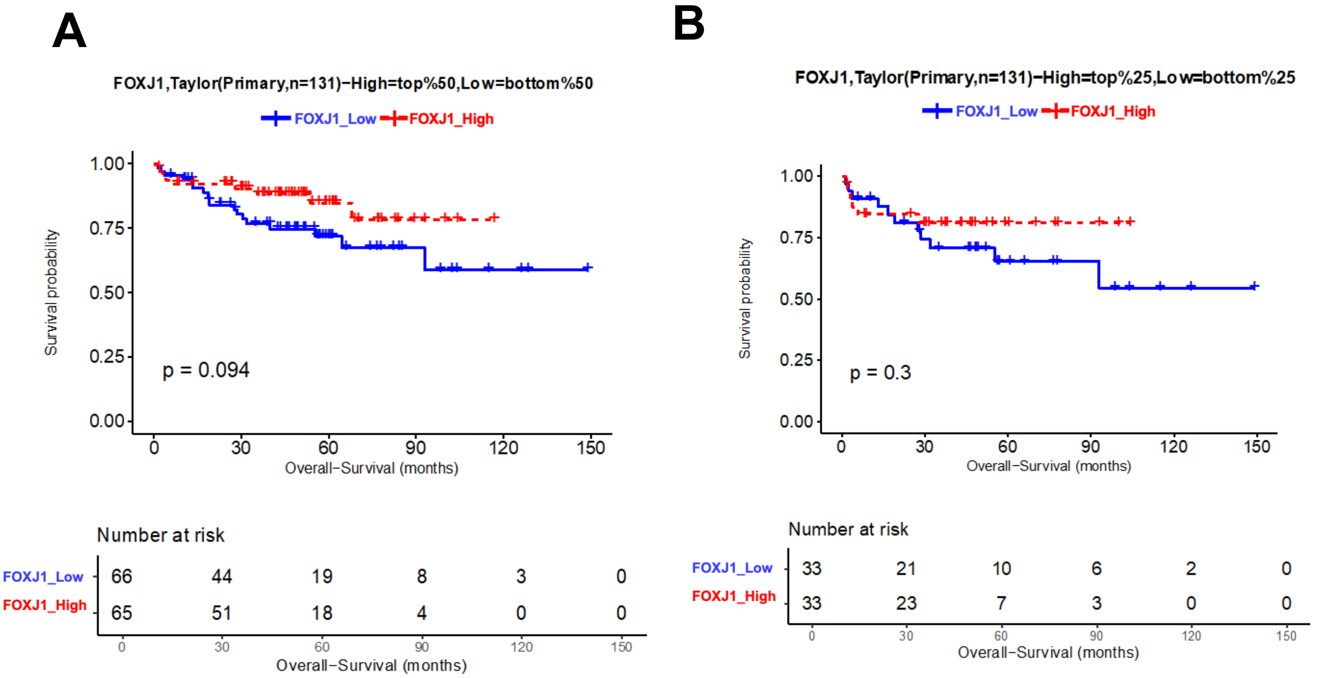


**Figure S18. Kaplan-Meier curves illustrating OS in patients from the Taylor PRAD dataset with high versus low level of FOXJ1. (A)** OS in top versus bottom 50% FOXJ1. **(B)** OS in upper versus lower quartile FOXJ1 expression.

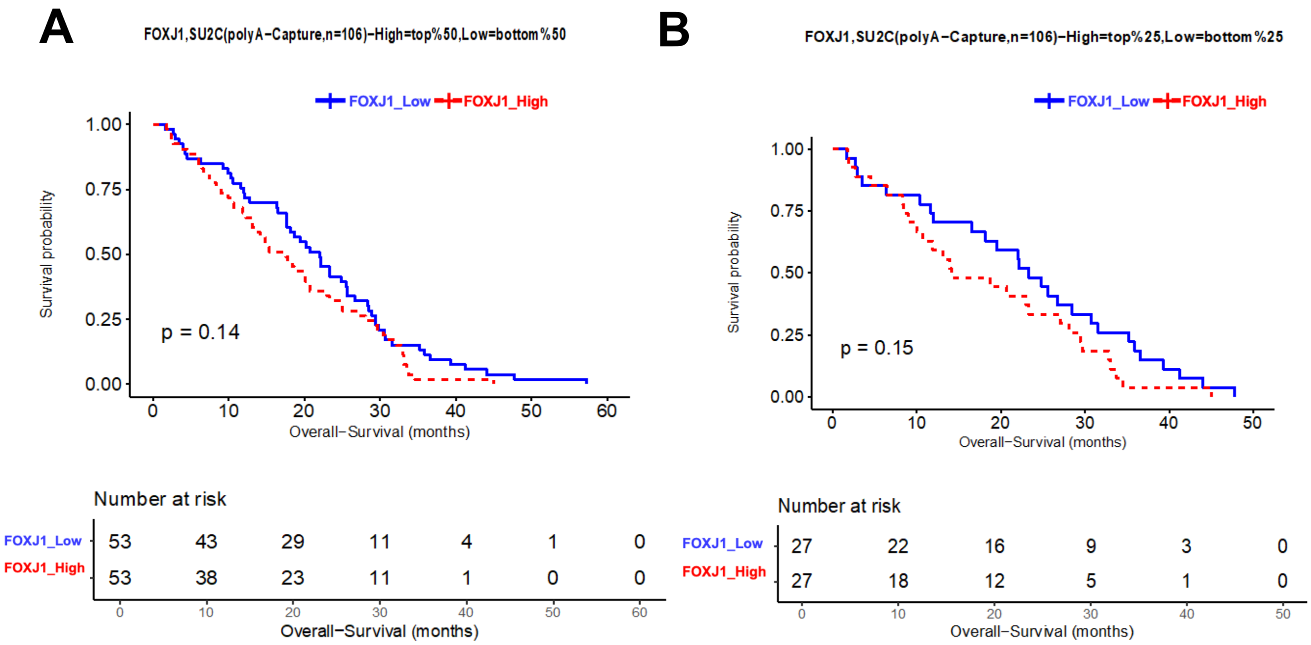


**Figure S19. Kaplan-Meier curves illustrating OS in patients from the SU2C CRPC dataset with high versus low level of FOXJ1. (A)** OS in top versus bottom 50% FOXJ1. **(B)** OS in upper versus lower quartile FOXJ1 expression.

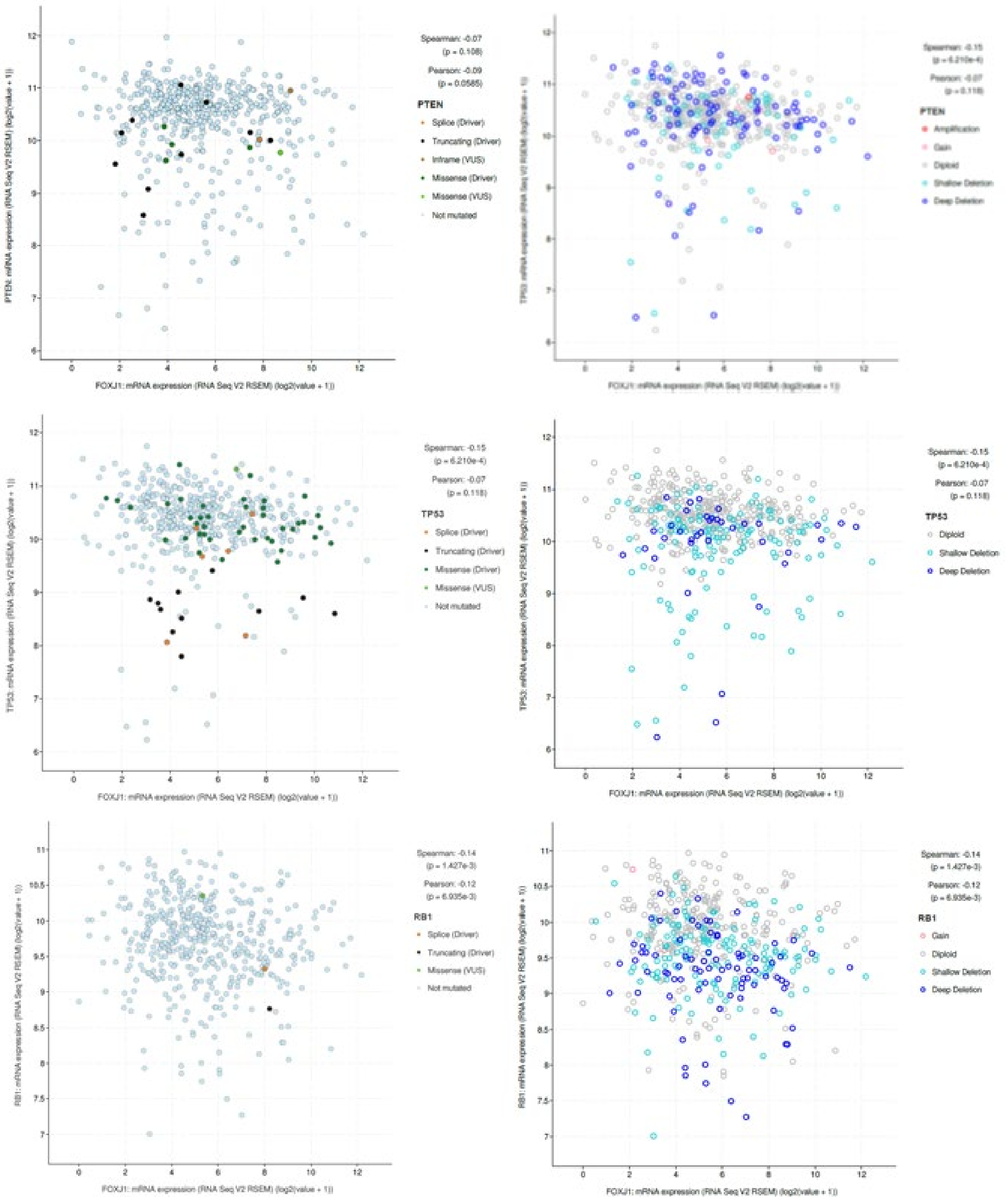


**Figure S20. Correlations between *FOXJ1* and *PTEN*, *TP53*, and *RB1* in primary PC.** Data on left panels shows mRNA levels and mutation status for *PTEN*, *TP53*, and *RB1*. Data on right panels show copy number alterations. Data are from the TCGA Firehose Legacy and were analyzed on cBioPortal.

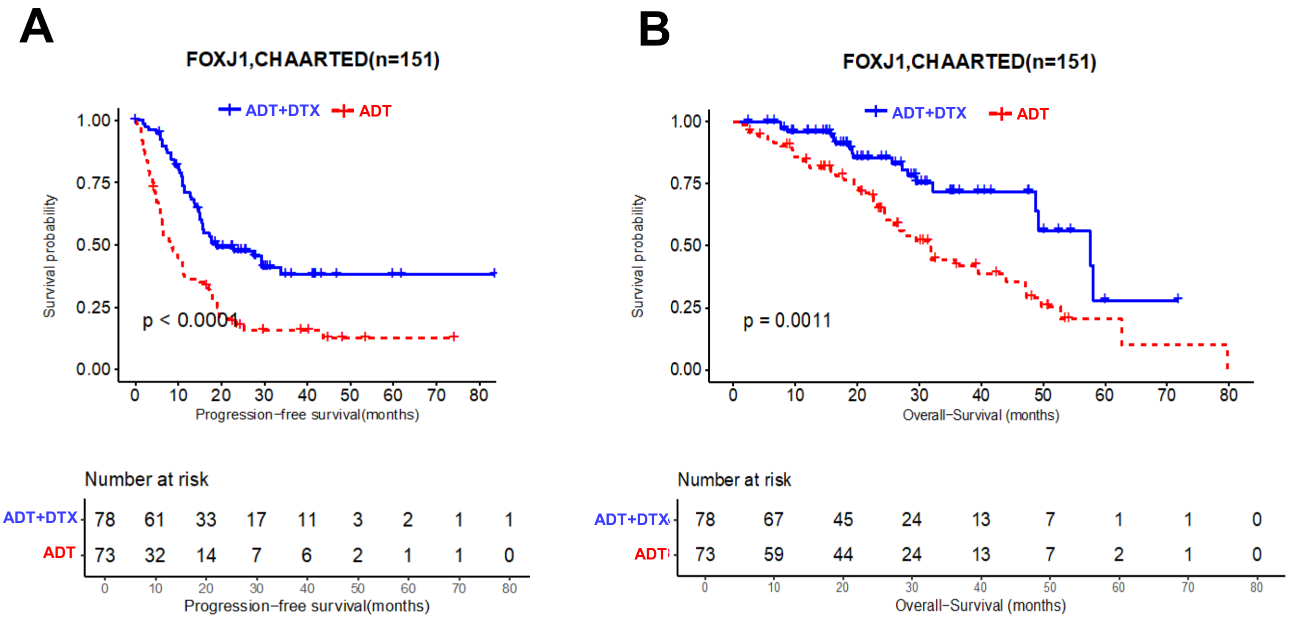


**Figure S21. Kaplan-Meier curves illustrating PFS and OS in patients from CHAARTED trial who were assessed for FOXJ1 expression prior to initiation of therapy. (A)** Improved PFS in Arm A (ADT + docetaxel) versus Arm B (ADT). **(B)** Improved OS in Arm A versus Arm B.

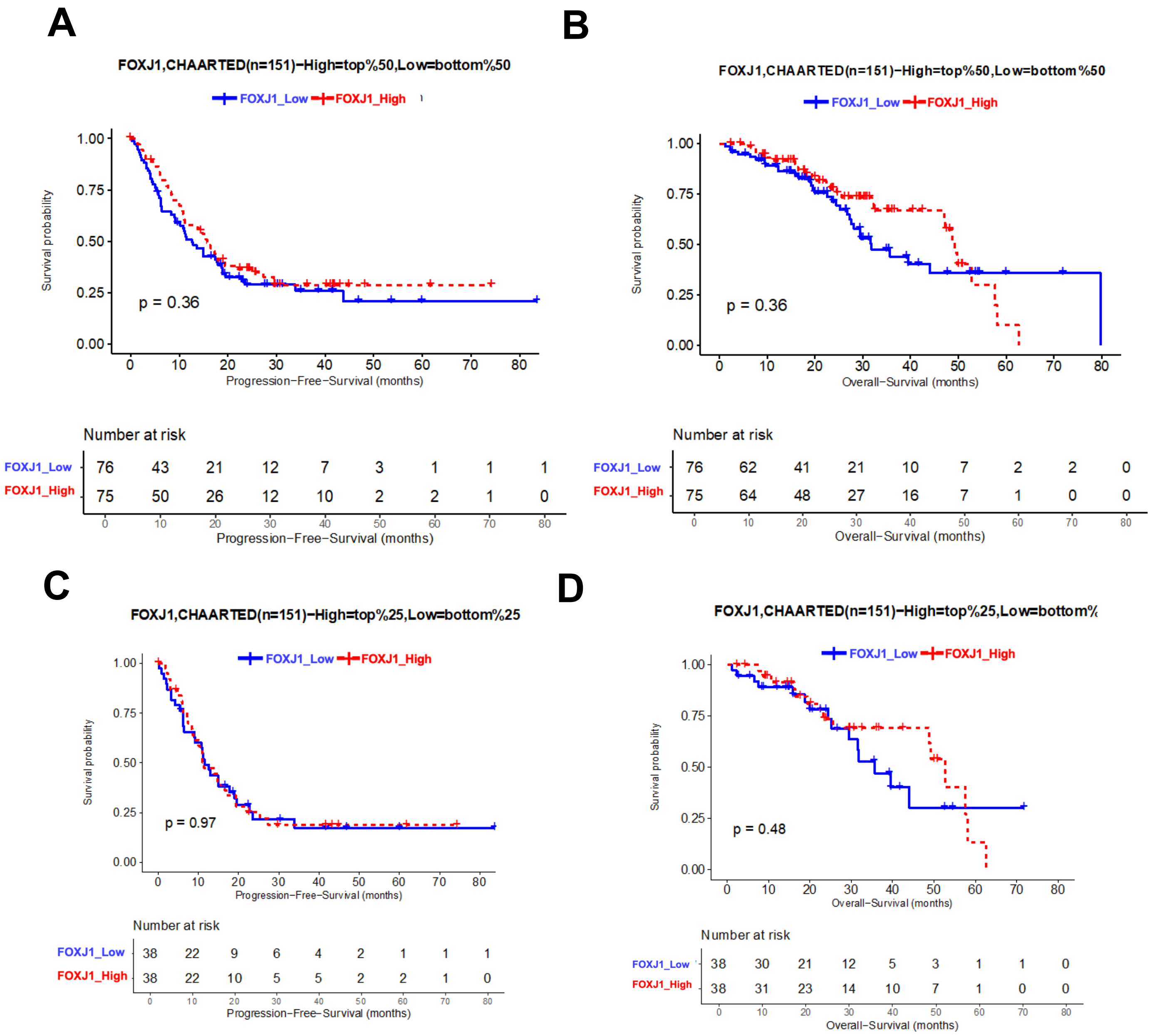


**Figure S22. Kaplan-Meier curves illustrating PFS and OS in CHAARTED patients with high versus low FOXJ1 expression. (A, B)** PFS (A) and OS (B) in top versus bottom 50% FOXJ1. **(C, D)** PFS (C) and OS (D) in upper versus lower quartile FOXJ1 expression.

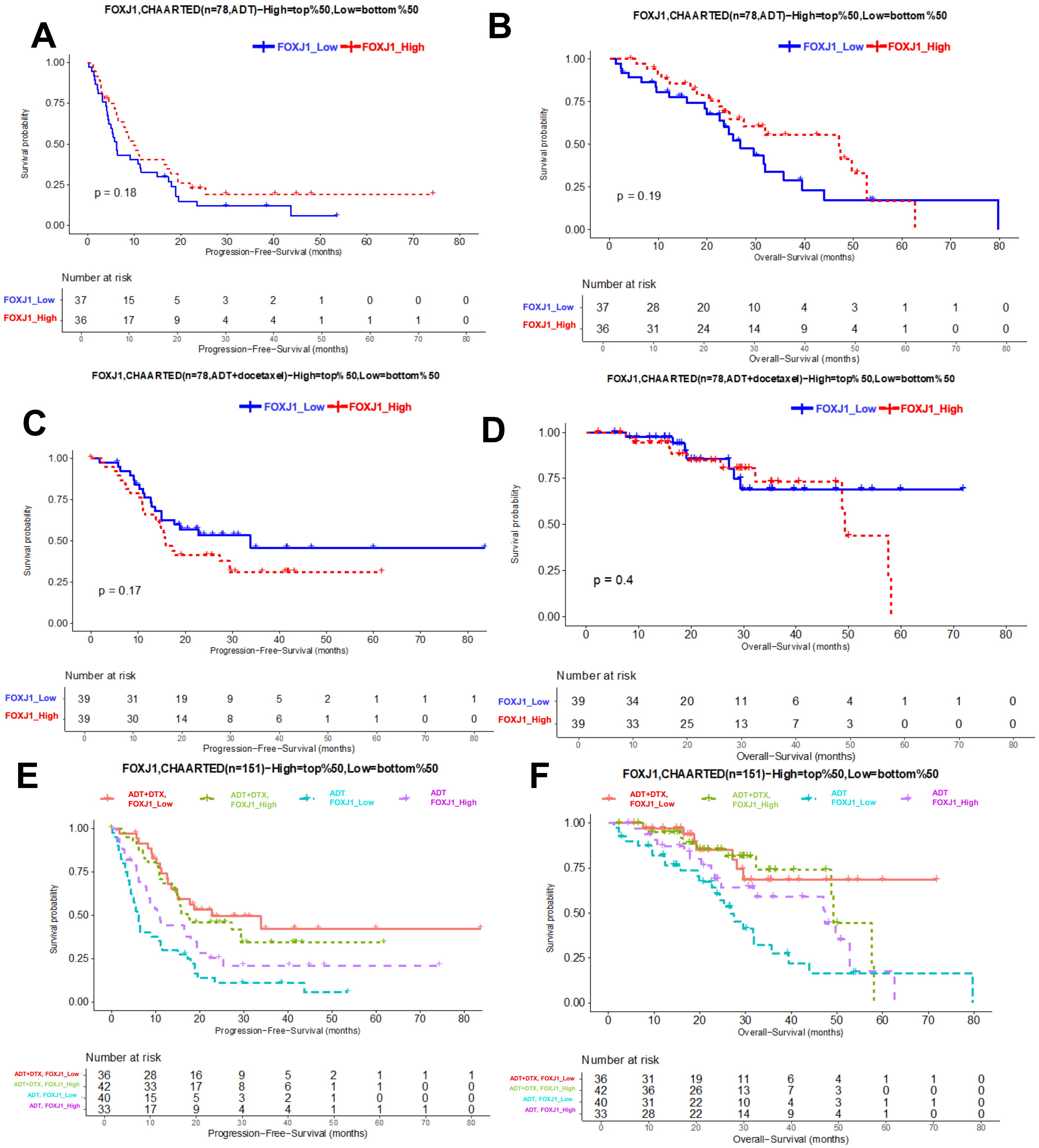


**Figure S23. Kaplan-Meier curves illustrating PFS and OS in CHAARTED patients with upper or lower 50% FOXJ1 expression treated on Arm A versus Arm B. (A, B)** PFS (A) and OS (B) in top versus bottom 50% FOXJ1 tumors treated on Arm B (ADT). **(C, D)** PFS (C) and OS (D) in top versus bottom 50% FOXJ1 tumors treated on Arm A (ADT + docetaxel). **(E,F)** PFS (E) and OS (F) in top and bottom 50% FOXJ1 tumors treated on Arm A versus Arm B.

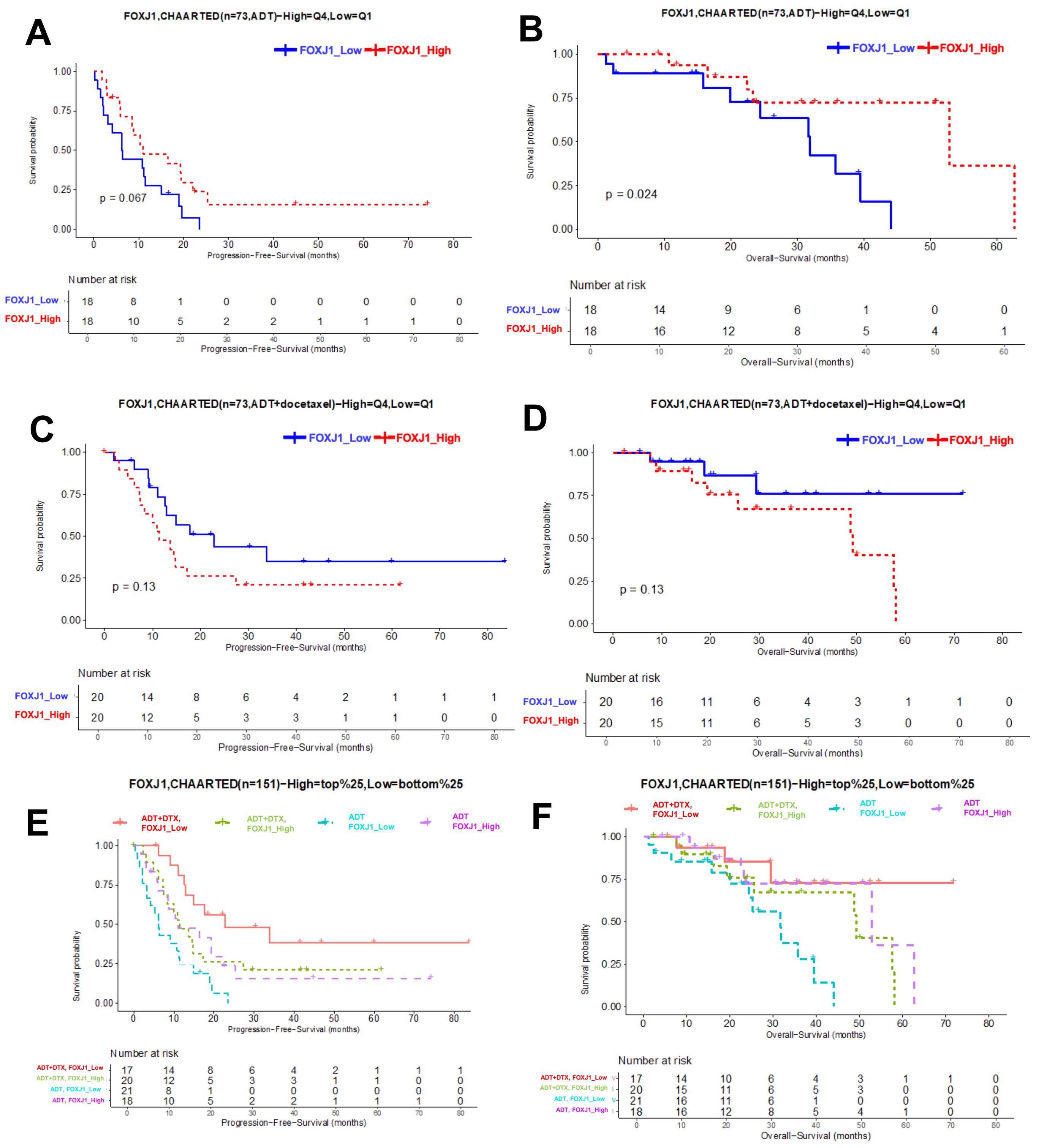


**Figure S24. Kaplan-Meier curves illustrating PFS and OS in CHAARTED patients with upper or lower quartile FOXJ1 expression treated on Arm A versus Arm B. (A, B)** PFS (A) and OS (B) in top versus bottom quartile FOXJ1 tumors treated on Arm B (ADT). **(C, D)** PFS (C) and OS (D) in top versus bottom quartile FOXJ1 tumors treated on Arm A (ADT + docetaxel). **(E,F)** PFS (E) and OS (F) in top and bottom quartile FOXJ1 tumors treated on Arm A versus Arm B.

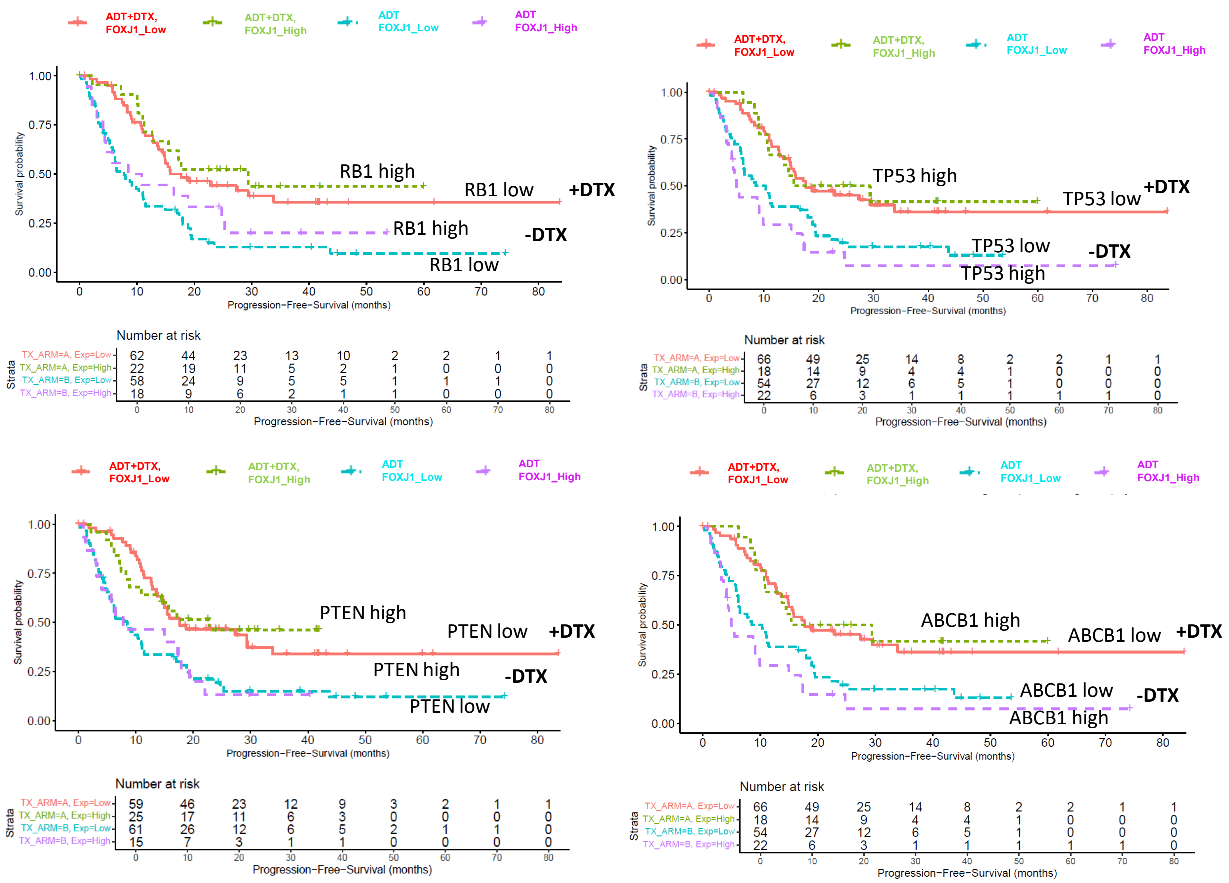

**Figure S25. Kaplan-Meier curves illustrating OS in CHAARTED patients based on expression of RB1, TP53, PTEN, and ABCB1.**

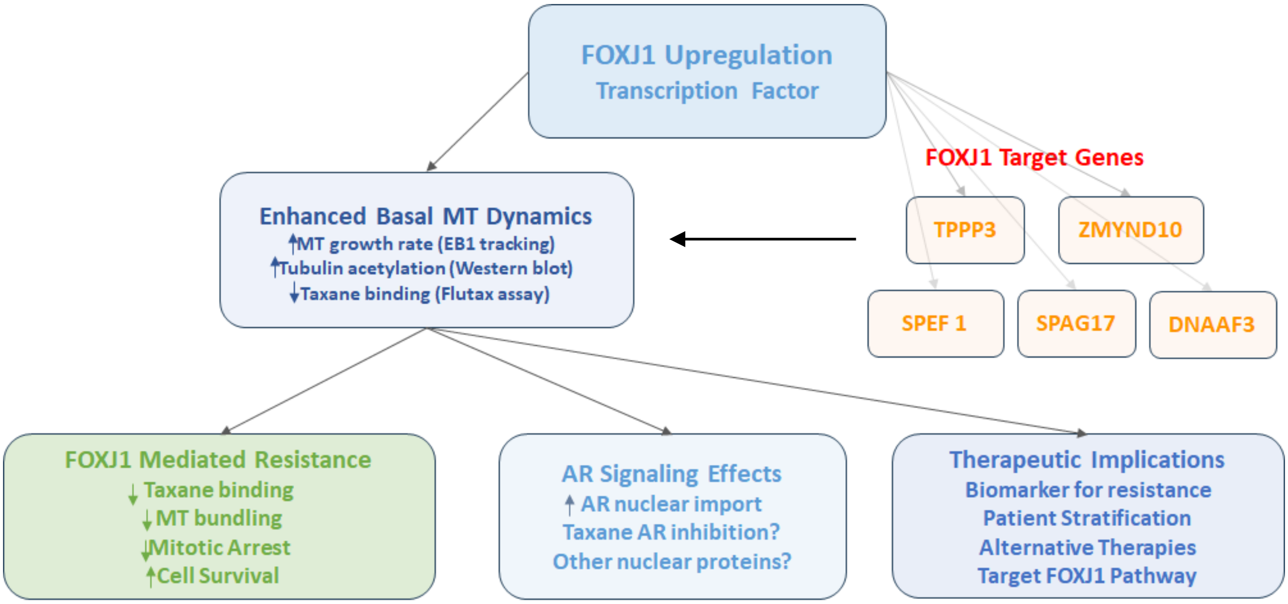


**Figure S26. Summary of FOXJ1 effects on MT dynamics and response to taxanes.**

### Supplementary Movies

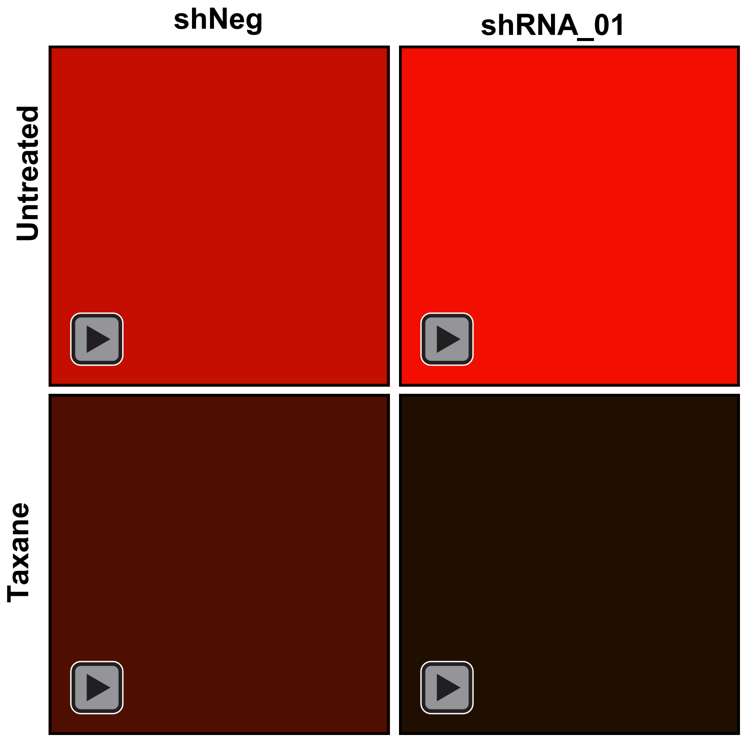


**Movie S1. Taxane treatment reduces the EB1 comet count.** Representative grayscale movies of DU145 shNeg and shRNA_01 cells, either untreated or treated with taxane, showing EB1 comets highlighted and tracked in purple dots over a 10-second timeframe using the Nikon Elements software. Taxane treatment significantly decreases the number of EB1 comets in both DU145 shNeg and shRNA_01 cells compared to untreated conditions.

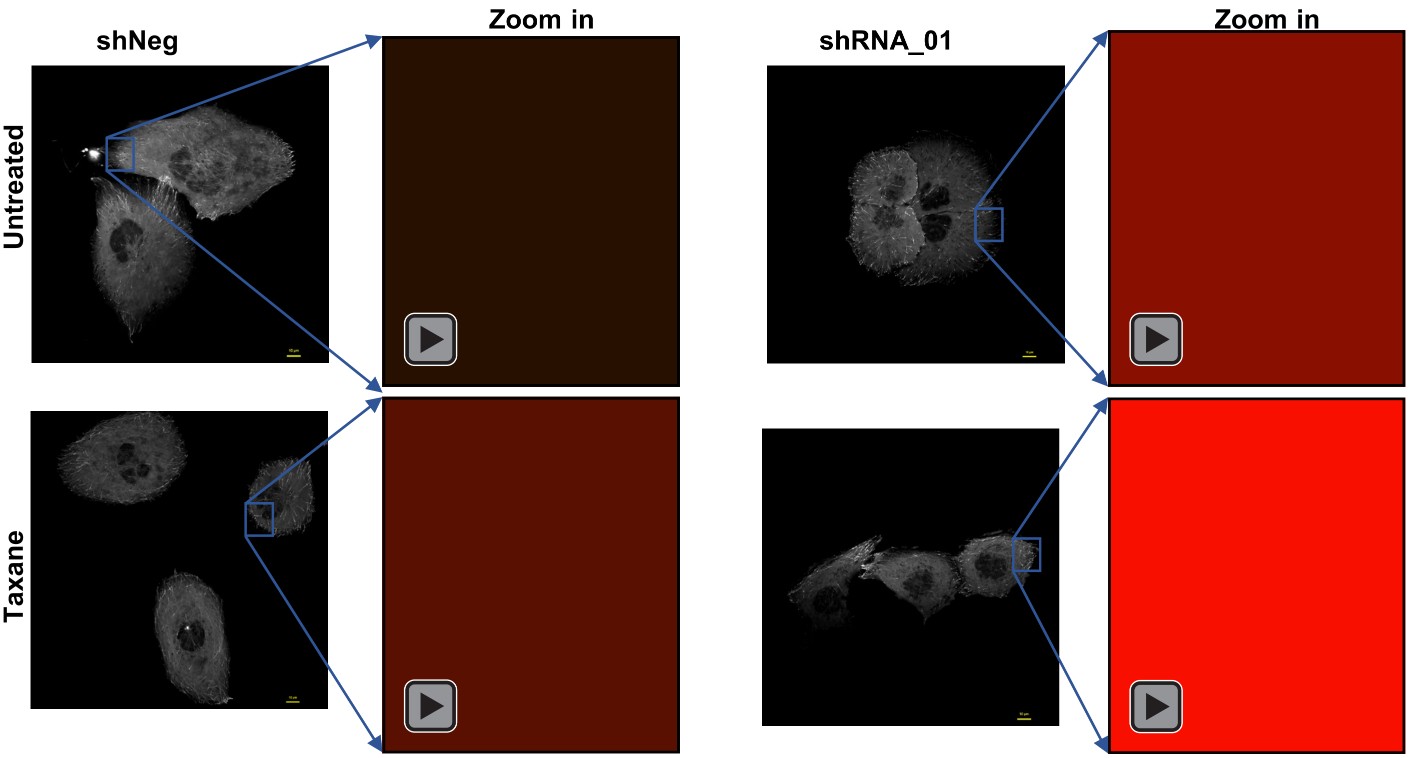


**Movie S2. FOXJ1 knockdown and taxane treatment significantly diminish EB1 comet speed.** Representative grayscale images of EB1-EGFP comets are shown for DU145 shNeg and shRNA_01 cells under both untreated and taxane-treated conditions, with a scale bar of 10 µm. The purple square indicates the region selected for detailed visualization. Zoomed-in movies of EB1-EGFP comets, tracked using NIKON Element software, show comet speeds over a 10-second timelapse, with a scale bar of 2 µm. These movies illustrate the dynamic behavior of EB1-EGFP comets, demonstrating a pronounced reduction in comet speed due to taxane treatment.

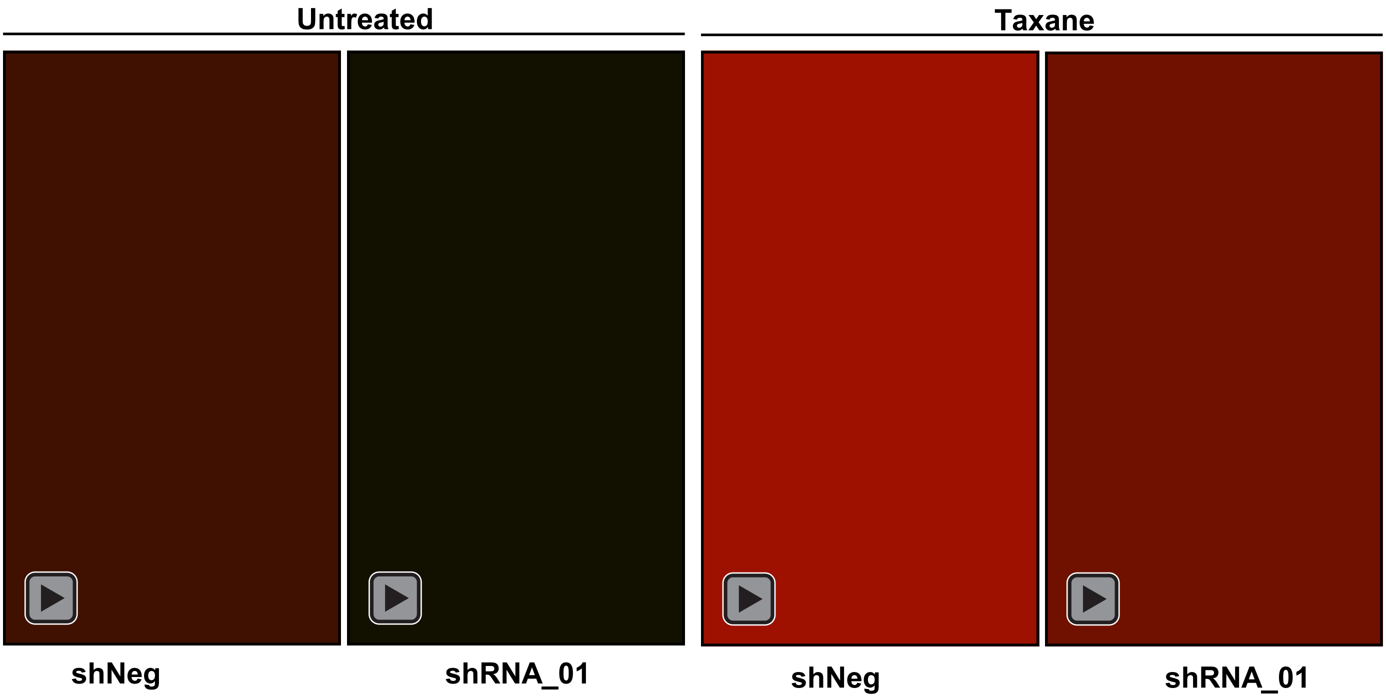


**Movie S3. Taxane treatment reduces EB1 comet trajectory length.** (Top) Representative time-lapse movies of untreated and taxane-treated DU145 shNeg and shRNA_01 cells were acquired to capture EB1 movement. Cells are shown in grayscale, while EB1 comets are displayed in pseudo-color using Nikon Elements software. Movies were recorded at 0.5-second intervals over 10 seconds. (Bottom) The time-course analysis depicts the trajectory of individual EB1 comets over 10 seconds. The x-axis represents time, and the y-axis shows the trajectory (path length) of EB1 comets as time progresses. In untreated shNeg cells, the maximum EB1 trajectory length is 12 µm, whereas in shRNA_01 cells, it is 9.5 µm. Following taxane treatment, the EB1 trajectory length decreases to 5 µm in shNeg cells and 4.5 µm in shRNA_01 cells.
