## Supplemental Videos for "FOXJ1 mediates taxane resistance through regulation of microtubule dynamics"

### Slide 1
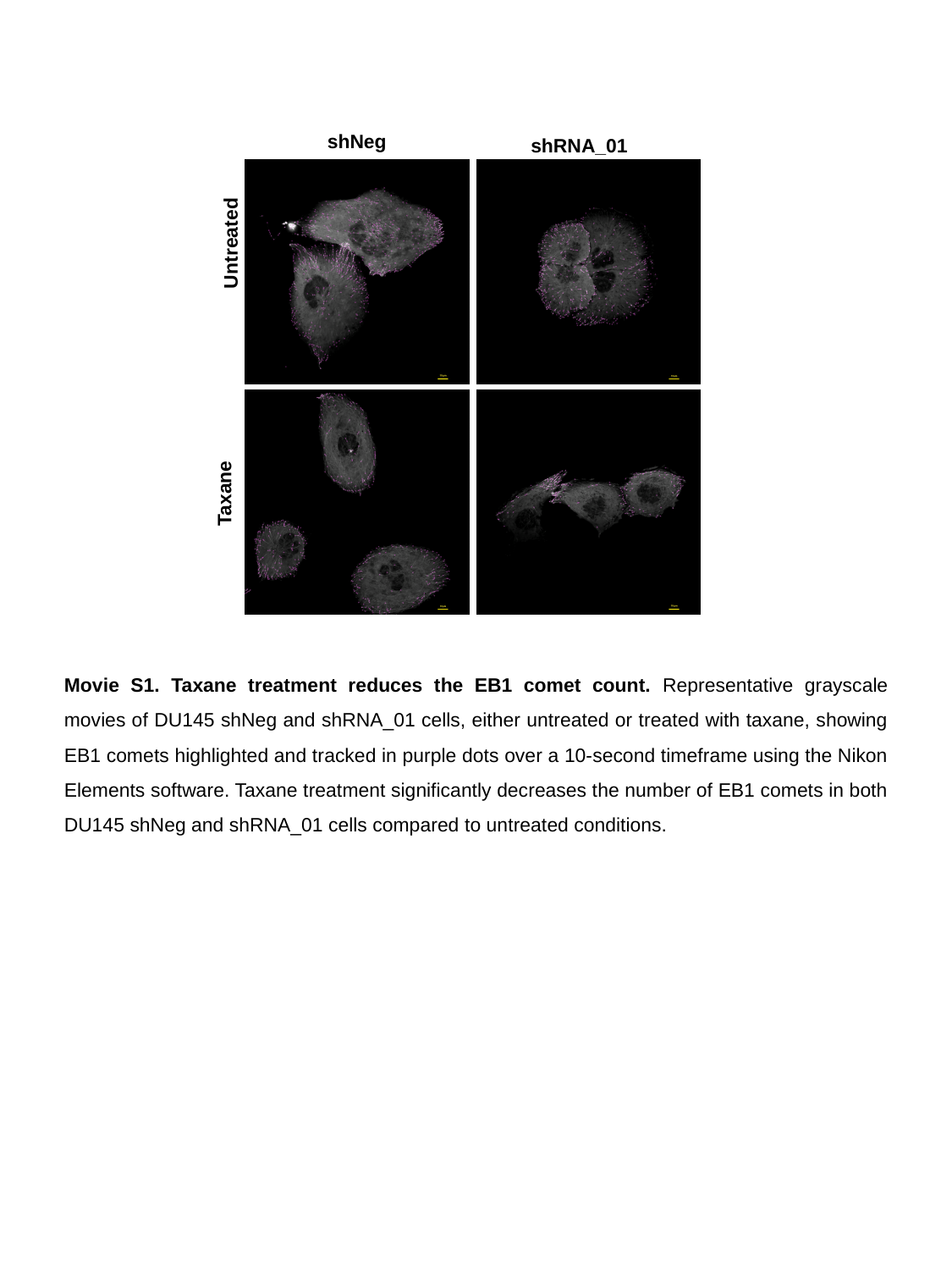

shNeg
shRNA_01
Untreated
Taxane
Movie S1. Taxane treatment reduces the EB1 comet count. Representative grayscale movies of DU145 shNeg and shRNA_01 cells, either untreated or treated with taxane, showing EB1 comets highlighted and tracked in purple dots over a 10-second timeframe using the Nikon Elements software. Taxane treatment significantly decreases the number of EB1 comets in both DU145 shNeg and shRNA_01 cells compared to untreated conditions.

### Slide 2
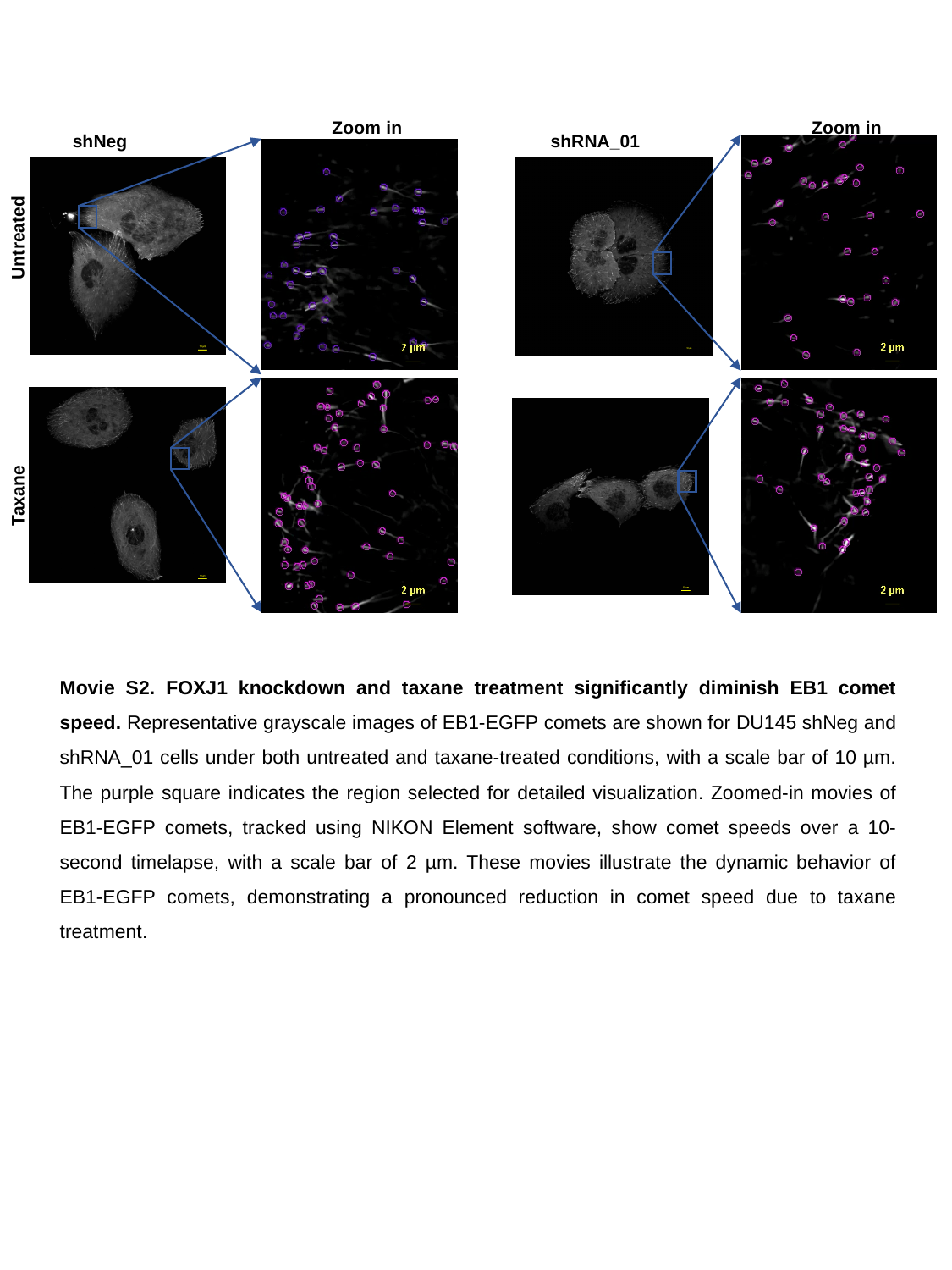

Zoom in
Zoom in
shNeg
shRNA_01
Untreated
Taxane
Movie S2. FOXJ1 knockdown and taxane treatment significantly diminish EB1 comet speed. Representative grayscale images of EB1-EGFP comets are shown for DU145 shNeg and shRNA_01 cells under both untreated and taxane-treated conditions, with a scale bar of 10 µm. The purple square indicates the region selected for detailed visualization. Zoomed-in movies of EB1-EGFP comets, tracked using NIKON Element software, show comet speeds over a 10-second timelapse, with a scale bar of 2 µm. These movies illustrate the dynamic behavior of EB1-EGFP comets, demonstrating a pronounced reduction in comet speed due to taxane treatment.

### Slide 3
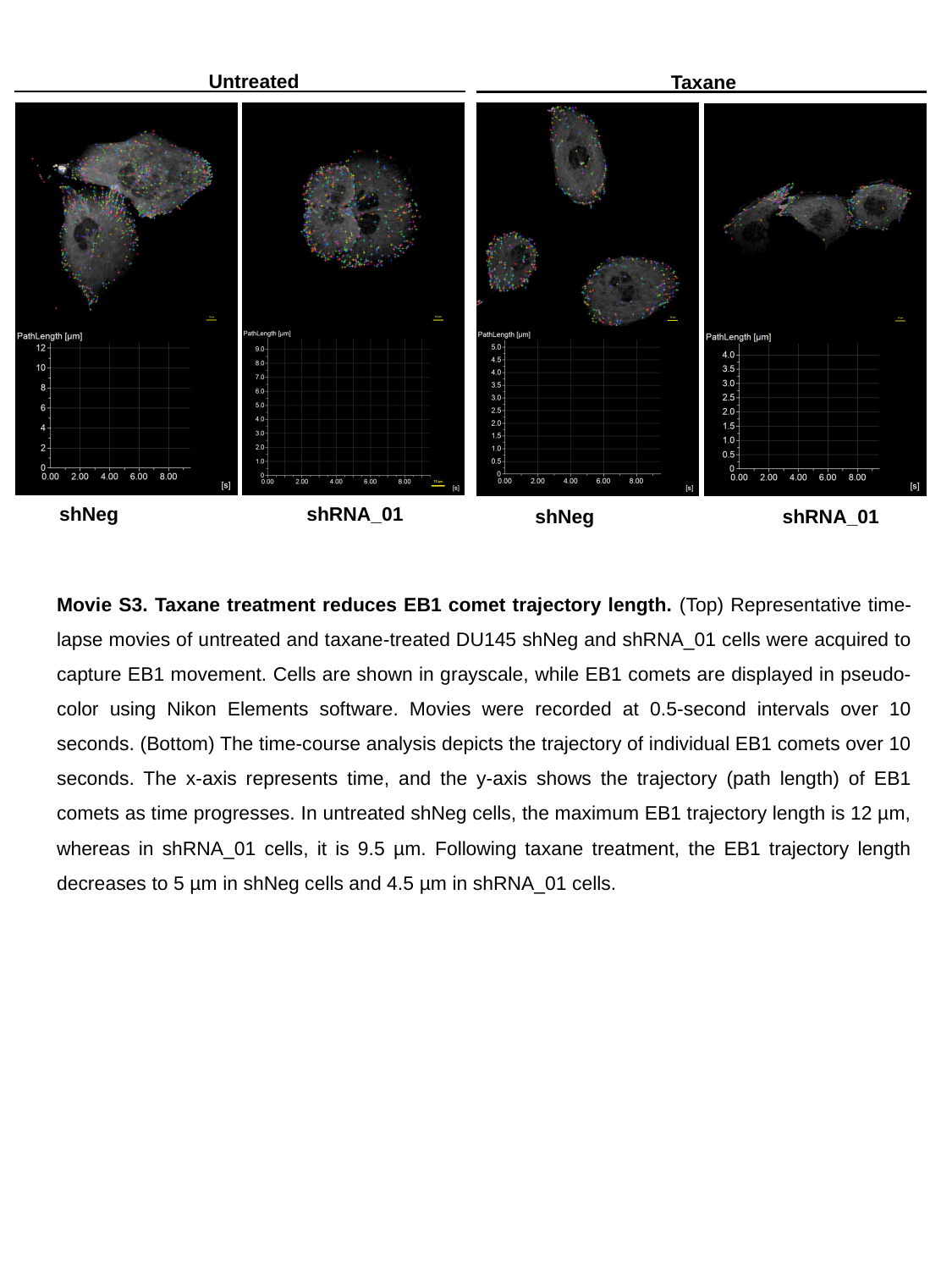

Untreated
Taxane
shRNA_01
shNeg
shRNA_01
shNeg
Movie S3. Taxane treatment reduces EB1 comet trajectory length. (Top) Representative time-lapse movies of untreated and taxane-treated DU145 shNeg and shRNA_01 cells were acquired to capture EB1 movement. Cells are shown in grayscale, while EB1 comets are displayed in pseudo-color using Nikon Elements software. Movies were recorded at 0.5-second intervals over 10 seconds. (Bottom) The time-course analysis depicts the trajectory of individual EB1 comets over 10 seconds. The x-axis represents time, and the y-axis shows the trajectory (path length) of EB1 comets as time progresses. In untreated shNeg cells, the maximum EB1 trajectory length is 12 µm, whereas in shRNA_01 cells, it is 9.5 µm. Following taxane treatment, the EB1 trajectory length decreases to 5 µm in shNeg cells and 4.5 µm in shRNA_01 cells.
